# Parameter-Efficient Fine-Tuning of a Supervised Regulatory Sequence Model

**DOI:** 10.1101/2025.05.26.656171

**Authors:** Han Yuan, Johannes Linder, David R Kelley

## Abstract

DNA sequence deep learning models accurately predict epigenetic and transcriptional profiles, enabling analysis of gene regulation and genetic variant effects. While large-scale training models like Enformer and Borzoi are trained on abundant data, they cannot cover all cell states and assays, necessitating new model training to analyze gene regulation in novel contexts. However, training models from scratch for new datasets is computationally expensive. In this study, we systematically evaluate a transfer learning framework based on parameter-efficient fine-tuning (PEFT) approaches for supervised regulatory sequence models. We focus on the recently published state-of-the-art model Borzoi and fine-tune to new RNA-seq datasets, enabling accurate and efficient analysis of gene regulation across different biological systems. Our results demonstrate that PEFT substantially improves memory and runtime efficiency while achieving high accuracy. The transferred models effectively predict held-out gene expression changes, identify regulatory drivers of differentially expressed genes, and predict cell-type-specific variant effects. Our findings underscore the potential of PEFT to enhance the utility of large-scale sequence models like Borzoi for broader application to study genetic variants and regulatory drivers in any functional genomics dataset. Code for efficient Borzoi transfer is available in the Baskerville codebase https://github.com/calico/baskerville.

## 2 Introduction

Deep learning models trained on DNA sequences accurately predict genomic profiles in unseen sequences across cell types and can be used to study regulatory drivers and genetic variant effects [1, 6, 24, 26]. Such models have been recently extended to predict RNA-seq coverage tracks [4, 19], allowing in-depth study of transcriptional, splicing, and polyadenylation regulation. Although these large-scale models have ample tissue, cell type, and assay representation during initial training, they nevertheless train on only a small proportion of datasets generated by the genome research community. Thus, many researchers could benefit from applying the predictive power of these models to their own data and biological systems of interest.

However, the optimal procedure for training models on new data to balance accuracy and computation remains unclear. Training from scratch is computationally expensive, and training on new data alone sacrifices the generalization benefits of multi-task learning across diverse assays and related cells. Transfer learning offers a promising alternative: pre-training models on large, diverse expression and epigenetic profiles, and then adapting this pre-trained model to specific experiments of interest [17, 23, 25, 27, 28, 29].

Traditional *fine-tuning* adapts a pre-trained model to new tasks by further training on domain-specific datasets, leveraging knowledge already encoded in the pre-trained model weights. However, as genomic models grow larger—with hundreds of millions of parameters—fine-tuning all weights becomes computationally prohibitive for many research groups. Parameter-efficient fine-tuning (PEFT) methods address this challenge by updating only a small fraction of model parameters while maintaining competitive performance. Originally developed for natural language processing and computer vision [8, 11, 12, 16, 18, 20, 33], PEFT approaches offer both computational efficiency and reduced overfitting risk, making them particularly attractive for adapting large genomic models to specialized datasets.

In this study, we implemented PEFT modules for both attention and convolutional layers in the Borzoi codebase and demonstrated their application to both bulk and single cell RNA-seq datasets that allow unique evaluation opportunities. For target datasets, we demonstrate strong gene expression prediction on unseen sequences, variant effect prediction, and the ability to recapitulate the epigenetic landscape and key regulatory motifs that drive transcriptional changes.

## 3 Results

### 3.1 Parameter-efficient fine-tuning improves transfer learning efficiency and performance

We used the recently published Borzoi deep multi-task model as our pre-trained foundation for transfer learning analyses [19]. This state-of-the-art model takes as input a 524 kb DNA sequence and applies convolution, transformer, and U-Net upsampling blocks (with 186 million parameters) to predict experimental aligned read coverage for 3,979 expression (CAGE, RNA) and 6,240 epigenetic (DNase, ATAC, ChIP) profiles at 32 bp resolution (Fig.1a). To enable more rapid experiments and benchmarking, we also trained a smaller version of Borzoi, referred to as Borzoi-lite, which takes a 393 kb input sequence, reduces model size throughout to 46 million parameters, and is trained on a more focused set of 4,754 human and mouse CAGE, RNA, DNase and ATAC profiles (Supplementary Table S1, Methods).

**Figure 1:**
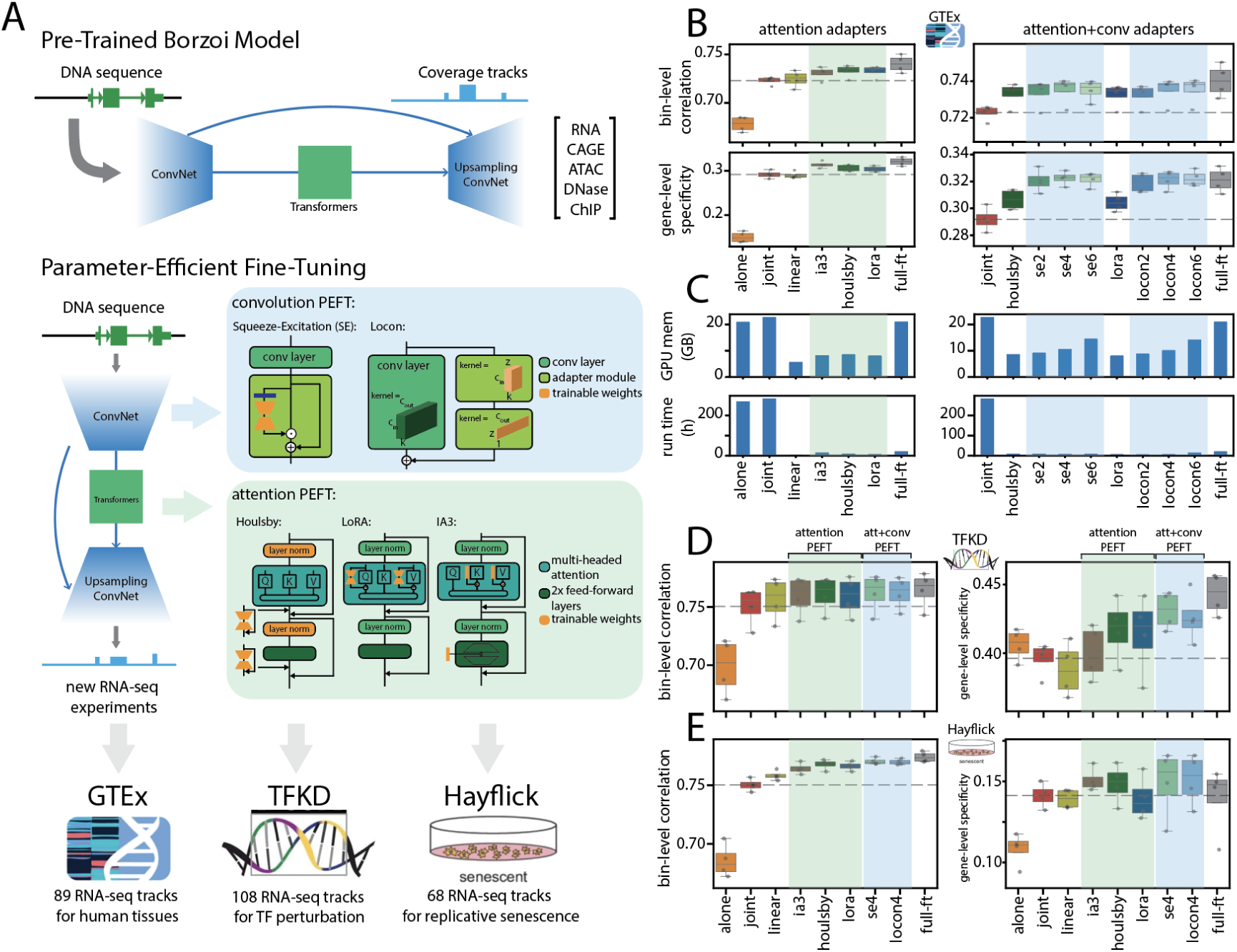
Borzoi parameter-efficient fine-tuning for RNA-seq transfer learning. (A) Borzoi is pretrained on a diverse set of functional genomics aligned read coverage tracks from human and mouse. In parameter-efficient fine-tuning (PEFT), we freeze all pre-trained parameters and insert learnable adapter modules (Methods). Blue shading indicates convolution PEFT modules Locon and squeeze excitation (SE), and green shading indicates attention PEFT modules (Houlsby, LoRA and IA3). (B) Borzoi-lite (pre-trained without GTEx) transfer performance on GTEx, measured by bin-level correlation and gene-level specificity for 4 folds on test sequences (Methods). Left: We compared attention-based PEFT methods (IA3, Houlsby and LoRA) to training from scratch on new data only (alone), jointly with all pre-training data (joint), fine-tuning with linear probing (linear), and full fine-tuning (full-ft). Right: We added SE adapters to the last 2, 4, and 6 convolution layers before attention in addition to Houlsby attention adapters (SE2, SE4 and SE6), or we added Locon adapters to the last 2, 4 and 6 convolution layers before attention in addition to LoRA attention adapters (Locon2, Locon4 and Locon6). We compared against joint, Houlsby, LoRA and full-ft. (C) Borzoi-lite transfer efficiency on GTEx, measured by average peak GPU memory (GB) and total runtime (hours) across folds. (D-E) Borzoi-lite transfer performance on the ENCODE TFKD dataset (D) and Hayflick senescence dataset (E), measured by bin-level correlation and gene-level specificity on test sequences. dynamically rescale convolution feature maps through learned channel attention mechanisms [13]. LoRA, IA3, and Locon adapters offer an additional deployment advantage: they can be merged back into the original model weights after training, eliminating any inference overhead while preserving the adapted functionality (Methods). This makes them particularly attractive for applications where inference speed and memory are critical.

Transferring a pre-trained sequence model to a new dataset can employ various strategies that demand different levels of computational resources. The most basic strategy involves replacing the final dense layer that transforms sequence representations into dataset-specific predictions to fit the new task. Training on new data can then update either only this final dense layer (linear probing) or all parameters through full fine-tuning. Linear probing freezes most of the pre-trained model, assuming it already captures general feature representations that can be adapted with a simple linear model, making it efficient but often yielding suboptimal performance. Full fine-tuning allows adaptation throughout the entire architecture but is resource-intensive for large models. Parameter-efficient fine-tuning (PEFT) bridges these approaches by updating only a minimal set of carefully designed parameters while maintaining competitive performance [8, 11, 12, 16, 18, 20, 33]. PEFT typically freezes all pre-trained parameters while inserting small learnable adapter modules that adjust layer computations to benefit new tasks.

We benchmarked three established PEFT approaches for Borzoi’s attention blocks: Houlsby adapters, LoRA, and IA3. These methods differ in their architectural modifications and parameter efficiency. Houlsby adapters insert bottleneck-structured modules after both the attention and feed-forward layers within each transformer block [11]. LoRA takes a fundamentally different approach by modeling weight updates as low-rank matrix decompositions [12]. IA3 introduces learnable scaling vectors to modulate attention weights and feed-forward activations [20] (Fig.1a, Methods).

Recognizing that convolutional layers in deep genomic models learn key sequence features such as transcription factor binding motifs [14, 35], we extended PEFT to Borzoi’s convolutional layers through two approaches. Locon adapts the LoRA principle to convolutional layers by decomposing convolutional kernel weight updates into low-rank factors [33]. Squeeze-excitation (SE) adapters

We evaluated transfer performance on held-out test sequences for three target datasets: GTEx, ENCODE TF perturbations (TFKD), and WI-38 senescence (Hayflick) RNA-seq [5, 7, 30]. We used the Borzoi-lite framework to train and compare multiple transfer approaches. Fold splits were identical between pre-training and transfer stages to ensure test sequences were unseen in both stages.

We first transferred the Borzoi-lite models pre-trained with GTEx data held out (Borzoi-lite-no-gtex model, Methods) to the GTEx RNA-seq dataset (Fig.1b, Methods) [7]. We evaluated model performance by 32 bp bin-level correlation between predicted and observed coverage on test sequences. Despite the diverse tissue representation of GTEx, we observed that transfer learning with even simple linear probing outperformed training from scratch on GTEx alone (mean R=0.725 vs. 0.678) and matched training from scratch on the entire pre-training collection jointly with GTEx (R=0.723). This highlights the benefits of transfer learning from a diverse pre-training collection. Introducing more free parameters in efficient structures via attention-based PEFT outperformed linear probing (mean R=0.733, 0.732, 0.731 for Houlsby, LoRA and IA3 vs. 0.725 for linear). Furthermore, the models benefited from adding convolution adapters alongside the attention adapters; mean Pearson R across folds improved from 0.733 to 0.734, 0.735, 0.735 when adding 2, 4 and 6 squeeze-excitation (SE) adapters to the Houlsby models, and from 0.732 to 0.732, 0.735, 0.735 when adding 2, 4 and 6 Locon adapters to LoRA models.

Regulatory sequence models are frequently used to study gene expression changes between tissues, cell types and states. We observed the same trend as above when we compared gene-level specificity prediction, which quantifies the correlation between observed and predicted gene expression after normalizing and subtracting the mean across tissues (Methods). We observed that adding attention adapters improved specificity compared to joint training or linear probing (mean R=0.306, 0.304, 0.314 for Houlsby, LoRA and IA3 vs. 0.292, 0.290 for joint and linear). After adding convolution adapters, specificity further improved from 0.306 to 0.320, 0.322, 0.321 when adding 2, 4, 6 SE adapters to the Houlsby models, and from 0.304 to 0.319, 0.321, 0.322 when adding 2, 4, 6 Locon adapters to LoRA models, closing the gap with full fine-tuning (mean R=0.321).

PEFT methods are highly resource-efficient. For example, LoRA used only 0.5% parameters, 35.8% GPU memory and 2.6% runtime compared to joint training from scratch, and 0.5% parameters, 38.6% memory, and 34.9% runtime compared to full fine-tuning. Adding 4 Locon adapters to LoRA models introduced only an additional 0.1% parameters, 11.0% GPU memory, and negligible change in runtime (Fig.1c). Additional benchmarks on test performance and efficiency can be found in Supplementary Fig.S1 and Supplementary Table S2.

We reached the same conclusions when fine-tuning pre-trained Borzoi-lite models to the TFKD and Hayflick RNA-seq datasets: a combination of attention and convolution adapters achieved high performance while maintaining efficiency (Fig.1de, Supplementary Fig.S1, Supplementary Table S3-4). Among attention adapters, Houlsby and LoRA often achieved similar performance, slightly outperforming IA3 (mean bin-level Pearson R=0.733, 0.732, 0.731 for Houlsby, LoRA and IA3 in GTEx; 0.768, 0.766, 0.764 in Hayflick; 0.761, 0.759, 0.760 in TFKD). Adding convolutional adapters (Locon and SE) showed comparable results overall, with performance improving when 2 and 4 adapters were inserted into convolutional layers. Adapting more convolutional layers did not yield further performance gains and significantly increased memory usage due to the need to store large intermediate activations. Due to efficient training and high performance across tasks, we selected the model with LoRA and 4 Locon adapters (referred to as Locon4) as our final transfer learning method for subsequent analysis. Notably, the Locon4 model allowed us to transfer the full Borzoi model within the 24 GB memory of a single RTX 4090 GPU.

### 3.2 Transfer learning predicts tissue-specific variant effects

Transferring the Borzoi model to custom RNA-seq data allows us to predict variant effects on gene expression and study regulatory mechanisms in specific biological samples. First, we asked whether variant effect predictions derived from PEFT are as accurate as those of the original model. To this end, we fine-tuned the pre-trained Borzoi-lite-no-gtex model to GTEx tracks and evaluated PEFT methods against baseline approaches in two GTEx eQTL tasks: (1) binary classification of fine-mapped eQTLs versus matched negatives, and (2) correlation between the predicted effect size and the measured effect size of fine-mapped eQTLs (Fig.2a, Methods).

**Figure 2:**
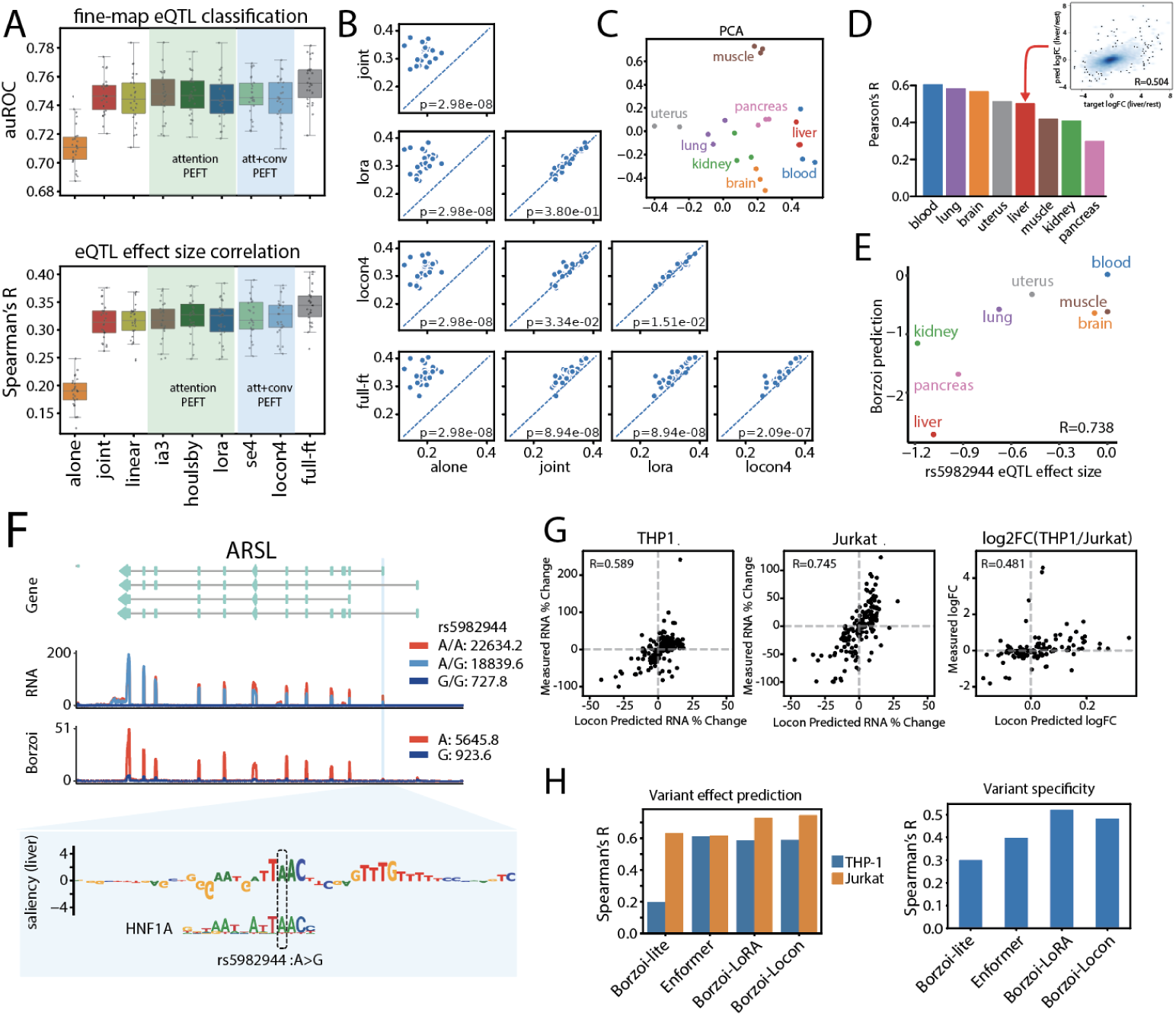
Transferred Borzoi models predict tissue-specific variant effects. **(A)** Borzoi-lite (pre-trained without GTEx) transfer performance for several transfer strategies on GTEx variant effect prediction, evaluated by classification auROC between fine-mapped eQTLs and matched negatives (top) and by correlation with measured effect size (bottom). **(B)** Comparison of eQTL effect size prediction across GTEx tissues, comparing models trained from scratch on GTEx alone, jointly with the pre-training dataset, or transferred with LoRA, Locon4, and full fine-tuning. Two-sided Wilcoxon sign-rank test p-values are indicated at the bottom right of each panel. **(C)** PCA plot of Locon4 model learned task representations from the final model layer parameters for eight representative tissues. **(D)** One-versus-rest log fold change (logFC) correlation between Borzoi-predicted and observed gene expression for test genes. A detailed correlation plot for liver is shown in the top-right corner. **(E)** Scatter plot of measured versus Borzoi-predicted eQTL effect sizes for rs5982944 across representative tissues. **(F)** From top to bottom: gene annotation of *ARSL*; GTEx RNA-seq counts averaged across individuals with A/A (red), A/G (light blue) and G/G (navy) alleles; Borzoi prediction for A and G alleles; Borzoi saliency scores around rs5982944 in liver. **(G)** Comparison of measured versus Locon4-predicted variant effects on *PPIF* expression in THP-1 (left) and Jurkat (middle) cells, and cell-type specificity (right, logFC difference between THP-1 and Jurkat). Spear-man’s correlation between predictions and measurements is shown in the top left of each panel. **(H)** Performance comparison of variant effect prediction for THP-1 and Jurkat, quantified by Spearman’s correlation between predicted and measured changes in *PPIF* expression (left), and cell-type specificity, quantified by Spearman’s correlation between predicted and measured logFC differences between THP-1 and Jurkat (right). Comparisons include Borzoi-lite models trained from scratch, Enformer CAGE tracks of matched cell types, Borzoi with LoRA transfer, and Borzoi with Locon4 transfer.

For the eQTL classification task, transfer learning approaches showed comparable performance to joint training from scratch on all pre-training data with GTEx, both outperforming training from scratch on GTEx alone (Fig.2a, top). For eQTL effect size prediction, the attention-based PEFT method LoRA slightly exceeded joint training from scratch (mean R=0.317 vs. 0.315 for LoRA and joint) and significantly outperformed training from scratch on GTEx alone (mean R=0.188). Adding convolutional adapters (Locon4) further improved performance over LoRA alone (mean R=0.322, *P* = 1.51 × 10^−2^). Full fine-tuning achieved the overall best performance on these tasks (mean auROC=0.756 for eQTL classification and mean R=0.343 for effect size prediction), likely benefiting from the diversity of the GTEx dataset (Fig.2ab, Supplementary Table S5-6).

We leveraged the GTEx data’s diverse tissue representation to evaluate whether transferred models learn tissue-specific regulation. Examining the final layer parameters of the transferred Locon4 model, we found that learned task representations accurately recapitulated known tissue relationships [34], with same-tissue tasks clustering together in principal component space (Fig.2c, Supplementary Fig.S2). This tissue-aware learning translated to strong performance predicting tissue-specific gene expression (mean R=0.488 predicting logFC across eight representative tissues, Fig.2d, Supplementary Fig.S2).

We observed that the Locon4 model also predicted tissue-specific variant effects. For example, rs5982944 (chrX:2964339:A:G) is a fine-mapped GTEx eQTL variant for *ARSL* in multiple tissues, with the strongest effects in kidney and liver. The transferred Borzoi accurately predicted the eQTL effect size across tissues (Fig.2e, Supplementary Fig.S2). In liver, the model predicted an almost complete loss of expression for the G allele, consistent with experimental measurements of individuals with this allele in GTEx (Fig.2f). Examining nucleotide saliency scores based on model gradients for *ARSL* prediction, we found that the variant disrupted the binding site of HNF1A, a known master regulator in kidney, liver, and pancreas, explaining the eQTL’s specificity (Fig.2f). This example highlights the power of transferred models to both learn and interpret tissue-specific variant effects.

To further evaluate performance on cell-type-specific variant effect prediction, we studied a set of short promoter insertions introduced in THP-1 and Jurkat cell lines to perturb *PPIF* expression, measured by Variant-FlowFish [21]. Here, we transferred the full-sized pre-trained Borzoi models to ATAC and RNA profiles from THP-1 and Jurkat cell lines, which took on average 10 hours on a single RTX 4090 GPU for LoRA/Locon4, compared to 279 hours when training Borzoi-lite models from scratch. On held-out test genes, Locon4 achieved the best prediction performance for each cell line (Pearson’s R=0.870, 0.862 for THP-1 and Jurkat, averaged across replicates), as well as cross-cell-type specificity (Pearson’s R = 0.331) (Supplementary Fig.S3).

For each model, we predicted the effect of each promoter variant on *PPIF* expression in both cell lines and compared our predictions to experimental measurements from variant-FlowFish (Methods). Locon4 achieved the best performance for within-cell effects (Spearman’s R=0.589, 0.745 for THP-1 and Jurkat), while LoRA slightly exceeded it for predicting cross-cell effects (Spearman’s R=0.521). Both techniques outperformed Borzoi-lite models trained from scratch and Enformer predictions derived from cell-matched CAGE (Fig. 2g, Supplementary Fig.S3). These analyses demonstrate that transferred models efficiently learn differential regulatory sequence factors across tissues and cells to enable accurate variant effect prediction.

### 3.3 Transfer learning reveals regulatory mechanisms of replicative senescence

The Hayflick dataset contains time-course bulk RNA and ATAC measurements of WI-38 cell lines either immortalized (IMT) or undergoing replicative senescence (SEN) [5]. We transferred the pre-trained full Borzoi model to RNA-seq data to evaluate its performance in predicting gene expression trajectories on held-out sequences, and in inferring senescence-associated transcription factor activity, validated using held-out ATAC-seq data.

Using Locon4, we successfully transferred the full Borzoi model using 15.6 GB of GPU memory and 54 training hours on a single RTX 4090, outperforming linear probing on test set performance (Supplementary Fig.S4). We observed that learned task representations from the final model layer parameters clearly separated the sample classes along the first principal component and revealed the progression of SEN cells toward replicative senescence (Fig.3a).

**Figure 3:**
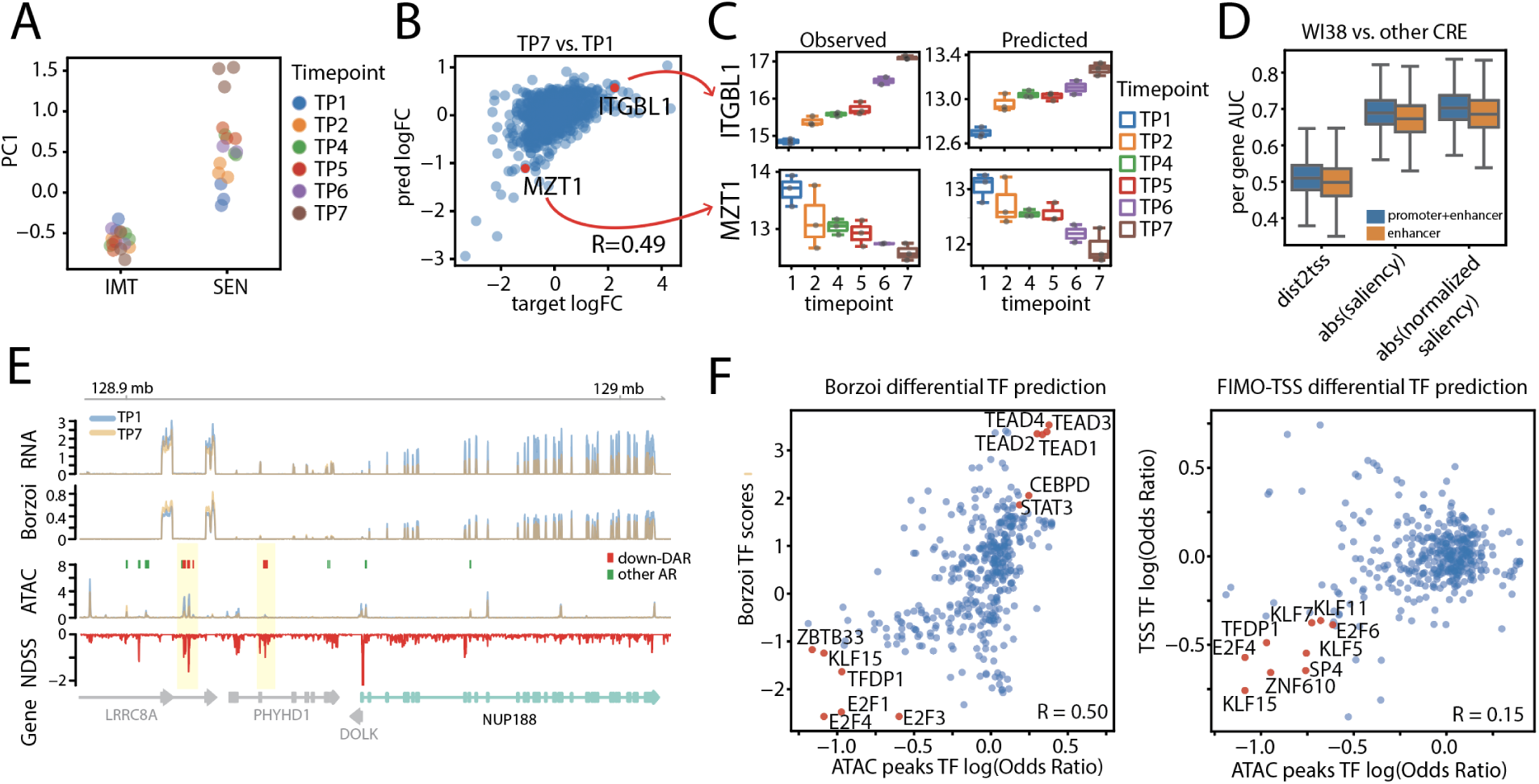
Transferred Borzoi models reveal the epigenetic landscape and key regulatory motifs of replicative senescence. **(A)** Distribution of the first principal component of sample embeddings derived from RNA-seq strand-concatenated final dense layer weights, showing separation between immortalized and senescent samples. **(B)** Scatterplot of predicted versus observed log fold change of senescent (SEN TP7) versus control samples (SEN TP1) on test genes. Two differentially expressed genes (*ITGBL1* and *MZT1* ) are labeled. **(C)** Observed versus predicted gene expression of *ITGBL1* (top) and *MZT1* (bottom) during replicative senescence (SEN TP7 vs. TP1). Y-axis shows log-quantile-normalized gene expression levels. **(D)** Performance comparison of different methods for identifying accessible WI-38 regulatory elements: TSS distance, absolute saliency, and normalized saliency. Blue boxplots show auROC distribution for all regulatory elements; orange boxplots show performance for enhancers only (>1kb from TSS). Normalized saliency accounts for regional activity through local smoothing and z-score normalization in a 10kb window (Methods). **(E)** Genomic tracks around senescence-repressed gene *NUP188* showing observed RNA-seq (TP1 control and TP7 senescent), Borzoi predictions, observed ATAC-seq with differentially accessible regions (red) and other accessible regions (green), and Normalized Differential Saliency Scores (NDSS) highlighting regulatory regions contributing to expression changes (Methods). **(F)** Left: RNA-derived Borzoi motif scores correlate with motif enrichment in DARs of matched ATAC-seq (Pearson’s R=0.50). Right: correlation between motif enrichment in TSS of DEGs and in DARs of matched ATAC-seq (Pearson’s R=0.15).

Focusing on SEN samples, we found that Borzoi predictions correlated with observed held-out gene expression changes during senescence (Pearson’s R=0.49 for TP7 versus TP1, Fig.3b, Supplementary Fig.S4). For example, the model accurately predicted the induction of *ITGBL1* and the repression of *MZT1* across time points, closely matching the observed expression patterns (Fig.3c). Visualizing the predicted and observed coverage tracks for example genes, we observed that Borzoi correctly predicted changes in coverage in the exonic regions for both senescence-induced and repressed genes (Supplementary Fig.S4).

In the baseline condition, gradient-based nucleotide saliency scores from the transferred Borzoi model identified key regulatory regions supported by WI-38 ATAC-seq. For genes of interest, we computed the saliency of aggregated TP1 RNA-seq coverage across gene exon bins with respect to the input nucleotides. To enhance signal detection in distal regions, we normalized the score by a regional aggregation statistic (Methods). The absolute value of the normalized gene saliency score prioritized WI-38 ATAC peaks over non-WI-38 peaks in the ENCODE cCRE atlas, outperforming unnormalized gene saliency or a simple distance-to-TSS baseline (Methods, Fig.3d). Comparing gradients between RNA-seq coverage tracks, we derived a Normalized Differential Saliency Score (NDSS) to highlight sequences contributing to differential gene expression regulation (Methods, Fig.3e, more examples in Supplementary Fig.S4)).

To identify TF motifs responsible for differential gene regulation during senescence (SEN TP7 versus TP1), we used the JASPAR2022 CORE motifs database and focused on motif instances around differentially expressed genes (DEGs) (Methods). A motif instance is considered to contribute significantly to differential gene expression if there is high NDSS magnitude and high specificity at the motif, measured via correlation between gradients to the reference and alternative nucleotides and the motif position weight matrix (PWM). Summarizing motif instance scores across DEGs (Methods), we derived Borzoi-predicted TF activities for the senescent trajectory, which correlated well with motif enrichment scores derived from ATAC-seq (Pearson’s R=0.50, Fig.3f). This approach successfully recapitulated key senescence-regulating TFs, such as TEAD and CEBP, as highlighted by Chan et al. [5]. In contrast, promoter-based motif enrichment analysis primarily identified TFs that regulate senescence-repressed genes but overlooked senescence-induced ones (Pearson’s R=0.15, Fig.3f), likely because senescence-induced genes are more enhancer-driven.

### 3.4 Transferred models predict transcriptional responses to perturbation

Within the ENCODE collection, researchers performed 108 RNA-seq experiments profiling transcription regulator knockdowns by CRISPRi or RNAi in the K562 cell line [30] (Supplementary Table S7), which we refer to as the TFKD dataset. To study TF regulation learned by transferred models, we transferred the full pre-trained Borzoi models to this data using Locon4. Analyzing the data before model training, we noted varying knockdown efficiency of target proteins and correspondingly weaker effects for incomplete knockdowns. Thus, we grouped experiments into categories based on regulatory knockdown efficacy to study model behavior. Borzoi could most reliably predict gene expression changes in the strong knockdown group, with decreasing and more variable logFC accuracy as knockdown efficiency decreased (Fig.4ab). Comparing predicted versus observed knockdown effects across experiments, we observed that Borzoi reproduced known regulator-regulator correlation relationships (Fig.4c, Supplementary Fig.S5). For example, knock-down of CTCF and the cohesin subunit RAD21 resulted in similar effects on downstream gene expression due to their well-established collaboration in determining regulatory domain boundaries [9].

**Figure 4:**
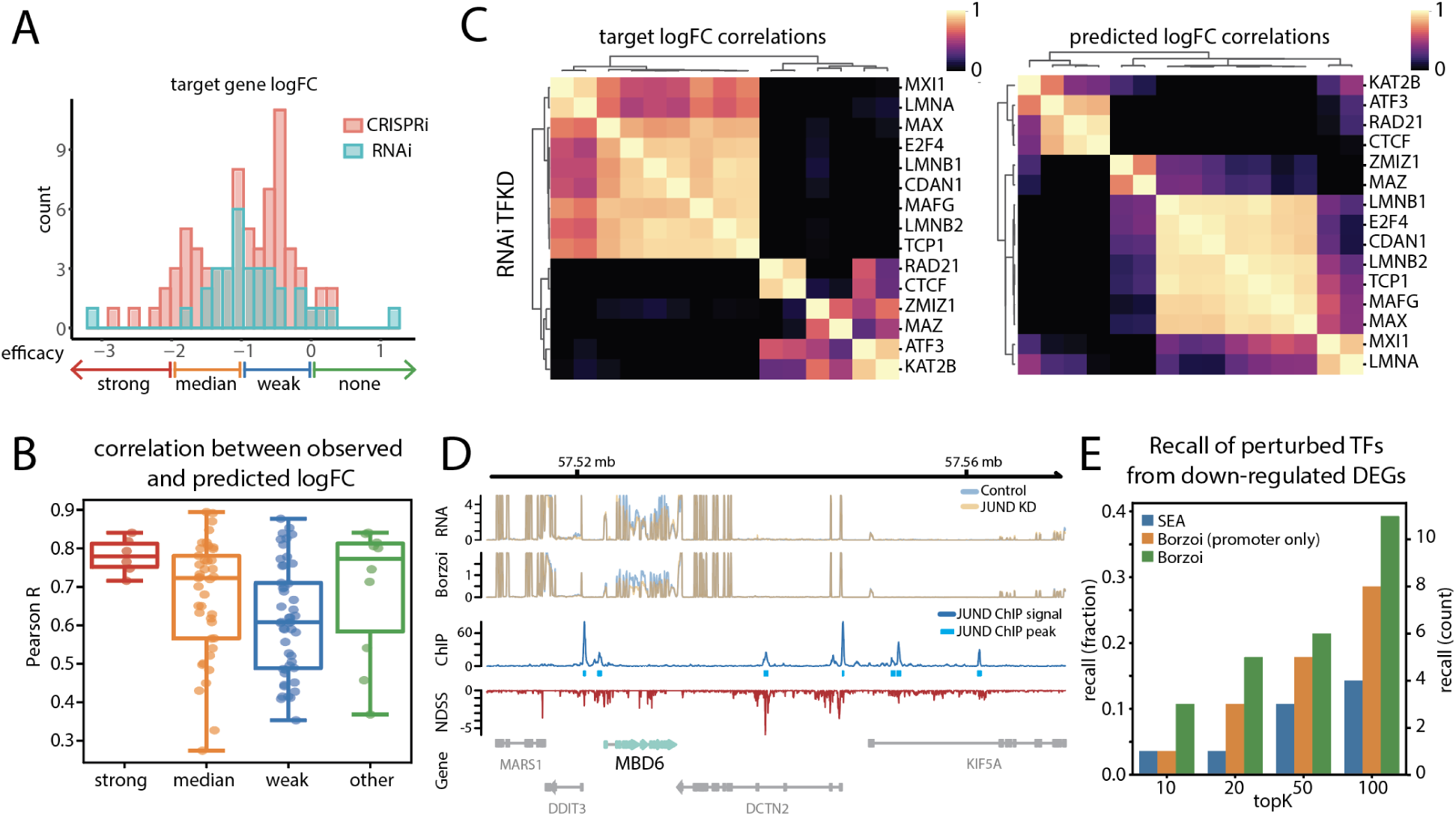
Transferred Borzoi models predict TF regulatory activity from perturbation RNA-seq. **(A)** Distribution of target gene logFC for 108 knockdown experiments. CRISPRi experiments are labeled in red and RNAi experiments in blue. Experiments with target gene logFC<-2, -2≤ logFC<-1, -1≤logFC<0, and 0≤logFC are categorized into strong, medium, weak effect and none groups. **(B)** Correlation between predicted and observed expression changes (quantile normalized, RNAi versus control logFC) on top 50% highly expressed test genes for each experiment, grouped by perturbation efficacy. **(C)** Correlation heatmaps of observed (left) and predicted (right) expression change (quantile normalized, RNAi versus control logFC) on test genes across RNAi experiments. (**D**) Predicted and observed tracks around a repressed gene *MBD6* after JUND CRISPRi knock-down. RNA: observed average RNA-seq tracks for control and JUND KD samples. Borzoi: Borzoi predicted RNA-seq tracks for control and JUND KD samples. ChIP: JUND ChIP-seq signal p-value track in K562. IDR thresholded peaks are shown in light blue. NDSS: Normalized Differential Saliency Scores (NDSS) comparing the saliency of *MBD6* between JUND KD and control sample (Methods). Gene: Gene annotation track. **(E)** For 28 knockdown experiments where the target TF has an existing motif in the JASPAR2022 database, we ranked all motifs by Borzoi activity score (using Borzoi scores, or Borzoi scores from promoter only), or by SEA enrichment score at promoters. We reported the recall as the fraction of experiments where the targeted TF is ranked among top K (k=10, 20, 50, 100).

By comparing saliency between perturbation and control gene expression predictions, we identified regulatory regions driving differential expression. For example, in the JUND knockdown, Borzoi NDSS revealed key regulatory regions associated with *MBD6* down-regulation, which corresponded to JUND-binding sites confirmed by ChIP-seq (Fig.4d). In Supplementary Fig.S5, we show another example of ATF3 knockdown effects on *TCL11L2*, where differentially salient regions contained ATF3 binding sites in the wild-type cells.

For the 28 perturbation experiments with strong or moderate efficacy and available JASPAR2022 motifs, we identified top down-regulated genes and evaluated whether Borzoi outperformed standard TSS motif enrichment (MEME Suite SEA) at identifying reduced activity of targeted TFs. Briefly, in a given TFKD experiment, we first identified significant motif instances contributing to down-regulated DEGs by Borzoi differential saliency, then calculated the overall TF contribution by aggregating scores across TF motif instances (Methods). When considering only promoter motifs, Borzoi NDSS better prioritized perturbed TFs compared to SEA motif enrichment scores. Relevant motifs highlighted by Borzoi NDSS in distal enhancer regions further improved recall (Fig.4e). In 5 of 28 perturbations, the targeted TF ranked among the top 20 based on Borzoi activity scores, whereas simple promoter motif enrichment (SEA) identified only one targeted TF across all experiments. For example, in the BACH1 knockdown experiment, Borzoi ranked BACH1 as the top TF driving down-regulated DEGs, while SEA ranked it at 257 (q-value = 0.456, Supplementary Fig.S5). Borzoi also allows examination of promoter versus enhancer motif contributions for each TF. For TFs such as BACH1, JUND, ATF3, and GATA1, enhancer scores were more predictive of their activity, whereas promoter scores were more relevant for known initiation factors like SP1 and NRF1 (Supplementary Fig.S5). These findings establish transferred Borzoi models as a powerful tool for dissecting transcriptional regulatory networks, particularly for identifying enhancer-driven regulation that conventional approaches fail to detect.

### 3.5 Transfer learning applied to single cell transcriptomic profiles

Even though Borzoi is pre-trained on bulk data only, researchers have explored transfer learning to cell-type-specific profiles derived from single cell genomics [10, 15]. To evaluate this capability with our workflow, we downloaded the PBMC 10x multiome (RNA+ATAC) dataset with 10k cells, clustered them into nine cell types, and generated pseudobulk profiles for each distinct cell type (Fig.5a, Methods). We then transferred the pre-trained Borzoi model to the scRNA pseudobulks using Locon4 on a single RTX 4090 GPU (Methods). We held out the scATAC data to validate TF activity prediction.

**Figure 5:**
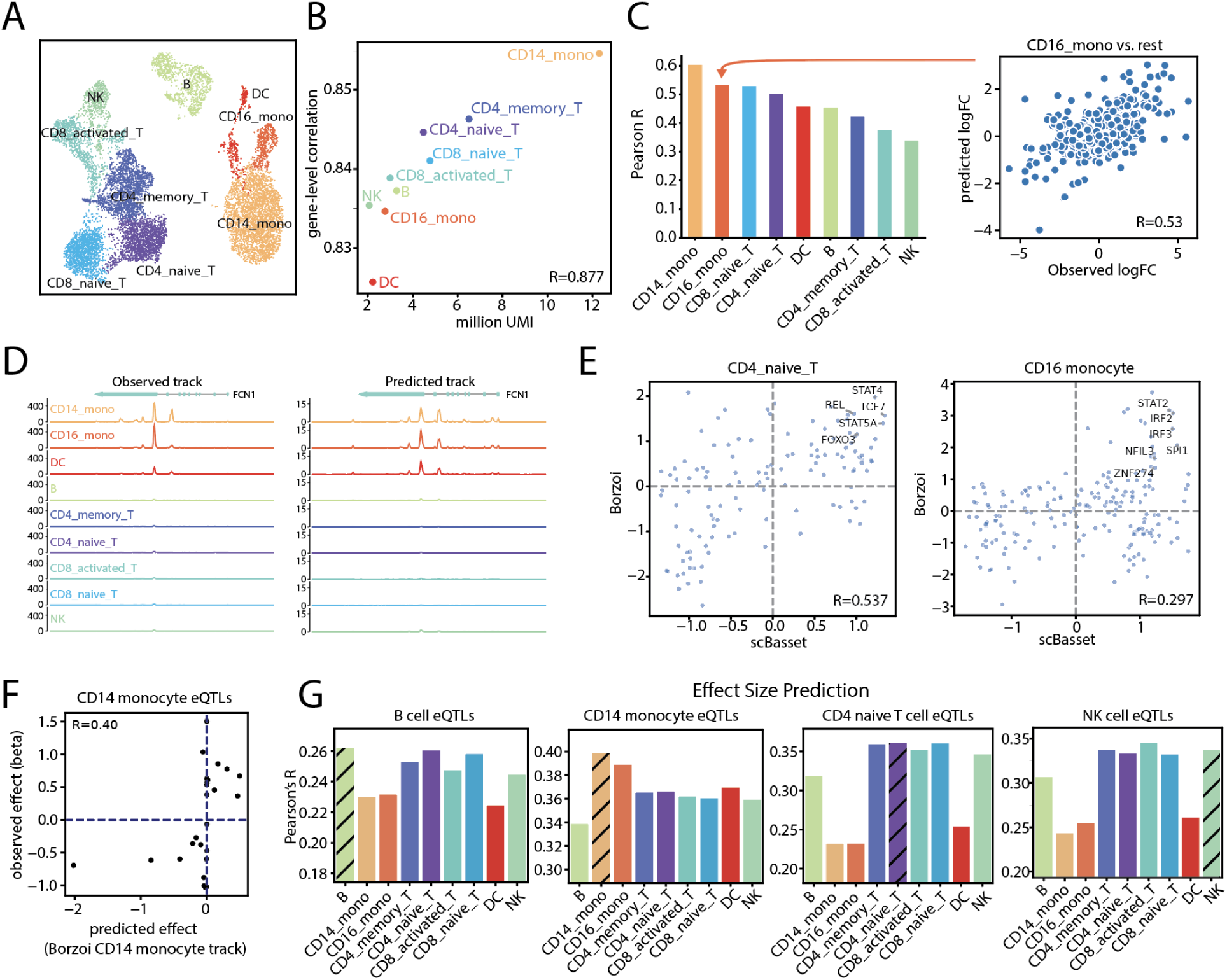
Transferred Borzoi models predict TF activity and variant effects in PBMC scRNA-seq. **(A)** UMAP visualization of PBMC cell embeddings with annotations. **(B)** Scatterplot showing the relationship between total UMI count and transfer performance, measured by gene-level correlation. **(C)** One-versus-rest log fold change (logFC) correlation between Borzoi-predicted and observed gene expression across test genes. A detailed correlation plot for CD16 monocyte is shown on the right. **(D)** Observed (left) and Borzoi-predicted (right) pseudobulk expression tracks for selected test gene *FCN1* across PBMC cell types. **(E)** Pearson’s correlation between TF activity inferred from scBasset using scATAC modality (x-axis) versus activity inferred from Borzoi using scRNA modality (y-axis) for CD4 naive T cells (left) and CD16 monocytes (right). **(F)**) Pearson correlation between measured and Borzoi-predicted variant effect sizes (log2FC) for fine-mapped eQTLs (PIP>0.9) in CD14 monocytes from the Onek1k cohort. **(G)** Pearson’s correlation between Borzoi cell-type-specific predictions and measured effect size of fine-mapped eQTL variants in B cells, CD14 monocytes, CD14 naive T cells, and NK cells.

On held-out test sequences, we observed lower bin-level correlation between prediction and observed coverage compared to typical bulk RNA-seq datasets (mean R=0.705, Supplementary Fig.S6), which likely reflects the domain shift introduced by 3′-biased poly-A capture and internal priming. When aggregated to the gene level, Borzoi achieved strong performance predicting gene expression counts and cell type specificity (Fig.5bc, Supplementary Fig.S6). As expected, performance correlated with UMI count in each pseudobulk profile (Fig.5b), but the spread between the minimum and maximum correlation was small (0.826-0.855).

Similar to our previous bulk RNA-seq analysis, we identified cell-type-specific regulatory TFs by scoring motif instances around marker genes using NDSS (Methods). Borzoi-inferred TF activity scores recapitulate known cell-type-specific TF regulators such as TCF7 in T cells and SPI1 in monocytes (Fig.5e). The scBasset method models ATAC peak status across single cells as a function of the underlying DNA sequence and delivers effective TF activity scores [34]. Comparing TF activity inferred from scRNA using Borzoi versus those inferred from scATAC using scBasset, we observed general concordance (mean R=0.258). Even in cell types where the methods diverged, they agreed on the top activating TFs. For example, the NK cell Borzoi RNA model may suffer somewhat from relatively fewer UMIs and lower specificity performance (Fig.5bc), but Borzoi and scBasset both highlighted major regulators RUNX2 and BCL11B (Supplementary Fig.S6).

To evaluate variant effect prediction performance, we applied the transferred model to fine-mapped, cell-type-specific eQTLs from the Onek1k cohort [32]. Across cell types, Borzoi predictions for the cell-type-specific log fold change effect size from the eQTL study were highly concordant (0.26-0.40 Spearman correlation, Fig.5f), and consistent with bulk metrics [19]. For all but one cell type, the matched cell predictions achieved the highest correlation among the other cell types, with the one outlier NK cells falling only 0.01 Spearman R behind the maximum. These results demonstrate that transferred Borzoi models successfully capture cell-type-specific regulatory mechanisms from single-cell data, achieving variant effect predictions comparable to those from matched bulk datasets.

## 4 Discussion

Our results demonstrate that parameter-efficient fine-tuning offers a practical solution for adapting large pre-trained regulatory sequence models to diverse transcriptomic datasets. By updating only a small fraction of model parameters, PEFT methods such as LoRA, Houlsby, IA3, Locon and SE achieve strong predictive performance with dramatically reduced computational cost compared to full fine-tuning. PEFT-adapted Borzoi models allow researchers to infer transcriptional regulation and predict tissue-specific variant effects using their own RNA-seq or scRNA-seq data as input, while operating on a single GPU with 20 GB of memory. These findings highlight PEFT as a lightweight and accessible strategy to extend state-of-the-art genomic models to new experimental settings.

A key practical consideration for PEFT deployment is determining the optimal placement and number of adapter modules. We systematically evaluated adapter placement across Borzoi’s convolutional layers, testing configurations with adapters in the final 2, 4, 6, or all convolutional layers preceding the attention blocks. Performance plateaued after 4 layers, while memory usage increased with deeper insertion due to the need to store large intermediate activations for gradient computation. This analysis revealed that while PEFT methods are parameter-efficient, memory overhead from backpropagation—not adapter parameters—becomes the practical bottleneck when adapting deep convolutional architectures.

We also found that full fine-tuning outperformed adapter-based methods in many cases, although its high GPU memory requirements may limit its practical application in some resource environments. This contrasts with findings in NLP, where adapter-based methods typically excel. We hypothesize that this difference arises because, in our RNA-seq transfer learning tasks, the target datasets contain the same number of examples (genome sequences) as the pre-training dataset, reducing the risk of overfitting.

The growing interest in applying large genomic models to single-cell data has sparked development of several complementary approaches. Recent work has demonstrated the versatility of the Borzoi architecture across different single-cell applications: Scooby adapts Borzoi to model continuous cell states at single-cell resolution, training on eight NVIDIA A40 GPUs [10]. Decima focuses on predicting gene-level expression matrices, performing full fine-tuning on one A100 GPU [15]. Our PEFT framework contributes to this ecosystem by prioritizing accessibility and flexibility. By maintaining Borzoi’s native track-level prediction capabilities while reducing computational requirements, our approach enables researchers with modest hardware resources to apply state-of-the-art sequence models to diverse genomic datasets. The ability to transfer effectively to bulk RNA-seq, single-cell, or multiome data using a single consumer-grade GPU democratizes access to these powerful modeling capabilities, potentially accelerating adoption across the broader genomics community.

For future work, combining adapter-based methods with techniques like quantization and distillation could further improve the efficiency of transfer learning. Including scRNA-seq data during the pre-training stage could also improve transfer performance to single cell datasets. Finally, introducing loss function terms that explicitly encourage accurate relative predictions between samples may boost the utility and interpretability of transferred models. In summary, our work demonstrates that PEFT enables efficient and accurate adaptation of large-scale regulatory sequence models to a wide range of transcriptomic datasets, unlocking their utility for diverse biological applications.

## 5 Methods

### 5.1 Pre-trained models

**Borzoi** takes as input a 524 kb DNA sequence and applies seven convolution blocks, eight attention blocks and two U-Net upsampling blocks to predict experimental aligned read coverage at 32 bp resolution for 7,611 human and 2,608 mouse genomic profiles, including RNA-seq, CAGE, DNase-seq, ATAC-seq, and ChIP-seq. Borzoi contains a total of 186 million parameters and was trained with a batch size of 2 across two A100 GPUs for 25 days (600 hours) until validation set accuracy plateaued. Four replicate Borzoi models were trained on the same train-validation-test splits. More detailed description can be found in the original Borzoi publication [19].

**Borzoi-lite** is a size-reduced version of Borzoi. It contains the same number of convolution, transformer and U-Net upsampling blocks as Borzoi. However, it takes as input a 393 kb DNA sequence, the number of filters in the convolutional layers is reduced by approximately half, and the number of heads in the transformer blocks is reduced by half. Borzoi-lite is trained to predict coverage at 32 bp resolution for 3,493 human and 1,261 mouse tracks containing CAGE, RNA-seq, DNase-seq, and ATAC-seq genomic profiles. Borzoi-lite contains a total of 46 million parameters and was trained with a batch size of 2 on one RTX4090 GPU for 13 days (312 hours) until validation set accuracy plateaued. For detailed comparison between Borzoi and Borzoi-lite see Supplementary Table S1. The genome was divided into six folds for cross-validation, where each model was trained on four splits and validated and tested on the remaining two. We pre-trained four Borzoi-lite models using folds 0, 1, 2, and 3 as the held-out test sets. We used Borzoi-lite as the pre-trained model for Hayflick and TFKD transfer benchmarks.

**Borzoi-lite-no-gtex** is a variant of Borzoi-lite which has the same architecture as Borzoi-lite and was pre-trained on the same data except for GTEx RNA-seq datasets. We used Borzoi-lite-no-gtex as the pre-trained model for GTEx transfer and variant effects benchmarks.

### 5.2 Transfer learning setup

We maintained the same train-validation-test split between the pre-training and transfer learning stages to prevent data leakage. For each Borzoi experiment, four pre-trained models were fine-tuned on the same train-validation-test split matching pre-training. For each Borzoi-lite experiment, four models were fine-tuned on the matching folds as pre-training.

Full fine-tuning and training from scratch with the full Borzoi model requires large GPU memory (>40GB) and cannot fit on a single RTX4090 GPU. Thus, for all benchmarks involving full fine-tuning and training from scratch, we used the Borzoi-lite architecture and Borzoi-lite/Borzoi-lite-no-gtex models as pre-trained models.

### 5.3 Parameter-efficient fine-tuning methods

#### 5.3.1 Attention PEFT: Houlsby

**Houlsby** is an adapter-based method which inserts learnable parameters into the transformer layers and freezes the remainder during fine-tuning [11]. Each adapter layer contains a dense layer projecting down to a low-rank space and a dense layer to project back, separated by a nonlinear activation. The adapter layer also contains a skip connection which allows it to approximate the identity function at initialization. We inserted two adapter layers in each transformer block before the residual connections as done in the original paper. During fine-tuning, all weights were frozen except for the adapters, layer normalization, and final head. We performed a hyperparameter search for the rank of the bottleneck *r* in {8, 16, 64} and learning rate in {1e-4, 1e-5}. The final hyperparameters chosen were: *r* = 8 and lr=1e-4 (Supplementary Fig.S7).

#### 5.3.2 Attention PEFT: LoRA

**Low-Rank Adaptation (LoRA)** hypothesizes that training updates to a dense weight matrix have low intrinsic rank during fine-tuning [12]. For a pre-trained weight matrix *W*_0_ ∈ ℝ*^d×k^*, Δ*W* refers to its cumulative gradient updates during fine-tuning. LoRA represents the updates with a low-rank decomposition 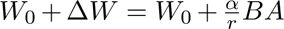, where *B* ∈ ℝ*^d×r^*, *A* ∈ ℝ*^r×k^*, the rank *r* ≪ min(*d, k*), and *α* is a scaling factor. During fine-tuning, the pre-trained weights *W*_0_ are frozen, while *A* and *B* are trainable parameters. For *h* = *W*_0_*x*, the modified forward pass yields:

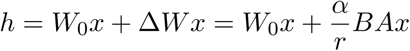

We implemented two modes of LoRA fine-tuning. In the default mode, we added adaptation only to the query and value weight matrices (*W_q_*, *W_v_*) as suggested in the original paper. In the full mode, we also added adaptation to the key and output matrices (*W_q_*, *W_v_*, *W_k_*, *W_o_*) of multi-head attention. During fine-tuning, all weights were frozen except for the LoRA modules and final head. We performed hyperparameter search for the rank *r* in {8, 16, 64}, scaling factor *α* in {4, 16}, mode in {default, full}, and learning rate in {1e-4, 1e-5}. Model performance was robust to hyperparameter choices, and the final hyperparameters chosen were: *r* = 8, *α* = 16, lr=1e-4, and mode=default (Supplementary Fig.S7). Because of the decomposition 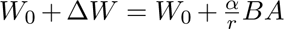, we can merge the weights of *BA* into *W*_0_ after LoRA fine-tuning to maintain the same architecture and computation as the pre-trained model, avoiding additional inference latency.

#### 5.3.3 Attention PEFT: IA3

**Infused Adapter by Inhibiting and Amplifying Inner Activation (IA3)** introduces learnable vectors that scale the attention and feed-forward activations in each transformer block [20]. For a transformer layer with query, key and value activations represented by *Q*, *K* and *V* , and feed-forward layer weights represented by *W*_1_ and *W*_2_, IA3 introduces learnable vectors *l_k_* ∈ ℝ*^dk^* , *l_v_* ∈ ℝ*^dv^* into the attention mechanism as:

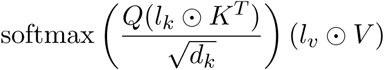

and introduces *l_ff_*∈ ℝ*^dff^* into the feed-forward network as (*l_ff_* ⊙ *γ*(*W*_1_*x*))*W*_2_, where *γ* is the feed-forward network activation and ⊙ represents element-wise multiplication with broadcasting notation (the (*i, j*)^th^ entry of *l* ⊙ *x* is *l_j_x_i,j_*) [20]. During fine-tuning, all weights were frozen except for the IA3 vectors and final head. We performed a hyperparameter search for the learning rate in {1e-4, 1e-5} and chose lr=1e-4 (Supplementary Fig.S7). Similar to LoRA, after IA3 fine-tuning, we can merge the vectors *l_k_*, *l_v_* and *l_ff_* into the original weights to maintain the same architecture and computation as the pre-trained model.

#### 5.3.4 Convolution PEFT: Locon

**Locon** is a variation of LoRA adapted to convolutional layers as introduced by Yeh et al [33]. Similar to LoRA, the cumulative gradient updates of a pre-trained weight matrix *W*_0_ are denoted as Δ*W* . For a 1-D convolutional layer, the weight update is denoted as Δ*W* ∈ ℝ*^Cout×Cin×k^* , where *k* represents the kernel size, *C_in_* represents the number of input channels, and *C_out_* represents the number of output channels. This matrix can be unrolled into 2D, represented as Δ*W* ∈ ℝ*^Cout×Cink^* and decomposed into Δ*W* = *BA* where *B* ∈ ℝ*^Cout×r^* and *A* ∈ ℝ*^r×Cink^* with rank *r*. Finally, the *B* and *A* matrices can be reshaped into *B* ∈ ℝ*^Cout×r×^*^1^ and *A* ∈ ℝ*^r×Cin×k^* . This is equivalent to two consecutive convolutions: first with weights *A*, input channels *C_in_*, output channels *r* and kernel size *k*; second with weights *B*, input channels *r*, output channels *C_out_* and kernel size 1. The modified forward pass for convolutional layer *h* = *W*_0_ ⊗ *x* yields:

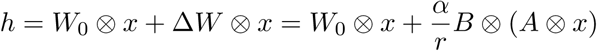

where *W*_0_, Δ*W* ∈ ℝ*^Cout×Cin×k^* , *B* ∈ ℝ*^Cout×r×^*^1^, *A* ∈ ℝ*^r×Cin×k^* , *r* is the rank, *α* is the scaling factor, and ⊗ represents the convolution operation.

We performed hyperparameter tuning for rank *r* in {4, 8} and *α* in {1, 16} and observed that performance was robust with respect to hyperparameter choice. Thus, we chose *r* = 4 and *α* = 1, which are the recommended defaults [33], and computationally more efficient. Similar to LoRA, Locon weights can be merged into the original architecture, avoiding inference latency. Throughout the manuscript, the **Locon** method indicates applying LoRA layers to all transformer layers, plus applying Locon to selected convolutional layers. Locon2, Locon4 and Locon6 indicate adding Locon to the last 2, last 4, and last 6 convolutional layers before attention, respectively. We combined Locon with LoRA for PEFT due to their similar operation and because both approaches allow for weight merging and result in no architecture change.

#### 5.3.5 Convolution PEFT: SE-adapter

We developed **SE-adapter** for convolutional layers inspired by the squeeze-and-excitation (SE) network [13]. Keeping all pre-trained weights fixed, we insert learnable SE blocks after convolutional layers and use them to fine-tune the convolutional activations. For an input *x* ∈ ℝ*^L×C^* of *L* sequence length and *C* channels, an SE block first applies global average pooling across the sequence length and outputs a 1D vector *h*_1_ ∈ ℝ^1^*^×C^*. The vector then undergoes a bottleneck two-layer feed-forward block followed by a tanh activation function to output a channel-scaling vector *h*_2_ ∈ ℝ^1^*^×C^*. The original input activation *x* is then re-scaled by *h*_2_ through channel-wise multiplication, and a skip connection is introduced to approximate the identity function at initialization, *h* = *h*_2_ ⊙ *x* +*x* where ⊙ represents channel-wise multiplication. The SE-adapter bottleneck rank was fixed at 16.

Throughout the manuscript, the **SE/Houlsby-se** method indicates inserting Houlsby adapters to all transformer layers, plus inserting SE-adapters to selected convolutional layers. se2, se4 and se6 indicate adding SE-adapter to the final 2, 4, and 6 convolutional layers before attention, respectively. We combined SE-adapter with Houlsby for PEFT because both approaches introduce architecture changes and result in additional inference computation.

### 5.4 Datasets

#### 5.4.1 RNA-seq data preprocessing

To account for the large dynamic range of RNA-seq data, Borzoi introduced a squashed scale transformation to the raw coverage tracks [19]. Here, we apply the same transformation to inputs of the Borzoi transfer models. The processing scripts accept BAM or BigWig files as input, and these are scaled so that one fragment is counted as a single event (with each position of the fragment contributing 1/fragment length). As a result, the total coverage across the genome sums to the read depth of the sample. The following preprocessing steps are then applied: (1) per-base coverage scores can be scaled by an optional factor, ‘scale’; (2) coverage scores are summed across 32-bp bins, followed by a square root transformation (sum stat=‘sum sqrt’); (3) if bin values exceed ‘clip soft’ after the square root, an additional square root is applied to the residual value; and (4) a final clipping step ensures values are capped at or below ‘clip’. For all our experiments, transformation was applied with parameters: sum stat=sum sqrt, clip=384, and clip soft=320. Custom scaling was applied for each dataset depending on input data type.

#### 5.4.2 GTEx

The GTEx dataset contains 89 unstranded RNA-seq coverage tracks from GTEx whole-tissue samples [7], uniformly processed by the recount3 project and later subsampled by Linder et al. [19, 31]. Briefly, for each recount3 meta-tissue, a subset of samples were included by k-means clustering the gene expression profiles with *k* = 3 (which collapsed to two clusters in several cases), and selecting one sample from each cluster. Transformation was applied with parameters: scale=0.02 to account for read length, sum stat=sum sqrt, clip=384, and clip soft=320.

#### 5.4.3 Hayflick

The Hayflick dataset aims to capture the replicative senescence process in WI-38 cells [5]. Bulk RNA and ATAC measurements were taken from wild-type WI-38 cells (SEN), which were intermittently sampled at 7 timepoints as they underwent replicative senescence. In addition to wild-type cells, hTERT-immortalized WI-38 cells (IMT) were grown in parallel and sampled as a control. This experiment contains RNA and ATAC measurements for two conditions (SEN and IMT) at 7 timepoints.

To transfer the Borzoi and Borzoi-lite models, we used only the RNA-seq tracks and held out the ATAC data for validation. Since the RNA-seq measurements are stranded, we separated the plus and minus strand reads from each experiment into two coverage profiles, leading to a total of 68 RNA-seq coverage tracks. For each RNA-seq track, aligned fragment counts were scaled by the inverse of their average length. Transformation was applied with parameters: scale=0.3, sum stat=sum sqrt, clip=384, and clip soft=320.

#### 5.4.4 TFKD

We refer to TFKD as the set of (mostly) transcription factor knockdown followed by RNA-seq experiments downloaded from ENCODE [30]. We downloaded BigWig files for 74 TF CRISPRi knockdowns in K562 followed by RNA-seq, and for 34 TF siRNA knockdowns in K562 followed by RNA-seq (also see Supplementary Table S7). For Borzoi-lite TFKD transfer learning, we transferred the pre-trained Borzoi-lite model to these 108 tracks. Downloaded ENCODE RNA-seq BigWig files are RPM normalized. Assuming a read length of 100 bp, the effective read count of an RPM normalized BigWig is 1*M* × 100 = 100*M* . We applied *scale* = 0.3 to normalize the dataset to an effective read depth of 100*M* × 0.3 = 30*M* , matching a typical RNA-seq experiment. Thus, transformation was applied with parameters: scale=0.3, sum stat=sum sqrt, clip=384, and clip soft=320.

For full Borzoi TFKD transfer learning, we transferred the pre-trained Borzoi model to 108 TFKD tracks, two K562 CRISPRi non-targeting control RNA-seq tracks, and two K562 siRNA non-targeting control RNA-seq tracks from the same batch. We also downloaded the corresponding gene quantification for these 112 experiments.

#### 5.4.5 Variant-FlowFISH

**Experimental design**: Variant-FlowFISH is an experimental technique used to measure the effects of CRISPR prime edits on gene expression for a single target gene in high throughput [21]. Martyn & Montgomery et al. used cell type-matched CAGE output of the Enformer model to rationally design a set of short indels for the *PPIF* promoter locus that would either minimize, maximize, or have no effect on *PPIF* expression in THP-1 or Jurkat cells (total n=164). A subset of the indels were designed to have no effect in either cell type (n=36), while the remainder of edits were designed to either maximize or minimize expression in THP-1 or Jurkat (n=128).

**Benchmark setup**: We used these Variant-FlowFISH measurements to benchmark different modeling approaches for predicting variant effects on gene expression. Specifically, we compared the performance of transfer learning (using PEFT methods on the full Borzoi model) against training smaller models from scratch and against pre-trained models without transfer learning. All models were evaluated on the same set of 164 designed variants for their ability to predict measured *PPIF* expression changes. For models requiring cell-type-specific training, we used matched ATAC-seq and RNA-seq data from THP-1 and Jurkat cell lines from the Engreitz Lab (ATAC-seq: GSM4706084, GSM4706085, GSM4706088, GSM4706089, GSM4706091, GSM4706092 [21]). A brief description of each model and how it was used in the benchmark comparison is given below.

##### Model implementations

*Enformer* : We used the THP-1 and Jurkat CAGE heads of the pre-trained Enformer model to predict variant effects in each respective cell line. Predicted *PPIF* expression was estimated as aggregated CAGE coverage in 3 bins overlapping the TSS, and variant effects were calculated as the log fold changes relative to reference sequence expression. Both reference and variant sequences were centered on the *PPIF* TSS, with predictions averaged across forward and reverse-complemented sequences and across sequences shifted by a few bp.

*Borzoi transfer learning (LoRA/Locon)*: We transferred the pre-trained Borzoi model to 6 RNA-seq and 6 ATAC-seq profiles from THP-1 and Jurkat cell lines using our PEFT methods. Variant scoring followed the same procedure as Enformer.

*Borzoi-lite*: We trained an ensemble of 4 Borzoi-lite model replicates (∼40 million parameters) from scratch on the cell-type-specific ATAC-seq and RNA-seq data. For variant scoring, we quantified predicted *PPIF* expression by aggregating coverage across the entire gene span from the corresponding RNA-seq output track. Reference and variant sequences were centered on the *PPIF* TSS. Predictions were averaged across forward and reverse-complemented sequences, sequences shifted by a few bp, and across the 4 model replicates.

#### 5.4.6 PBMC multiome

The 10x PBMC RNA+ATAC multiome dataset was downloaded from https://support.10xgenomics.com/single-cell-multiome-atac-gex/datasets/1.0.0/pbmc_granulocyte_sorted_10k. We followed the muon tutorial for preprocessing and cell type annotation using the scRNA modality https://muon-tutorials.readthedocs.io/en/latest/single-cell-rna-atac/pbmc10k/3-Multimodal-Omics-Data-Integration.html [3]. We then merged cell types pDC and mDC into DC cells, memory B and naive B into B cells, MAIT into CD8-activated T cells, and intermediate mono into CD14 monocytes based on marker gene expression to increase the depth of the pseudobulk profiles. We annotated a total of 9 cell types: B cells, CD14 monocytes, CD16 monocytes, CD4 naive T cells, CD4 memory T cells, CD8 naive T cells, CD8 activated T cells, DC, and NK cells.

We aggregated cells by cell type and generated stranded pseudobulk profiles, normalized to reads per million (RPM), in BigWig format. Given that the average read length in 10x scRNA-seq is 90 bp, each RPM-normalized BigWig file has an effective total read count of 1*M* × 90 = 90*M* , meaning the sum of values across the BigWig equals 90M. To match the pseudobulk UMI counts, we applied a scaling factor of 0.03 during Borzoi transfer, resulting in an effective read count of 90*M* × 0.03 = 2.7*M* , approximately the median UMI count in pseudobulk profiles.

We then transferred the full pre-trained Borzoi model to the nine scRNA-seq pseudobulk profiles using Locon4, yielding a total of 18 tracks (accounting for strandedness).

### 5.5 Transfer learning performance metrics

#### Bin-level correlation

For each track, we computed the correlation between predicted and observed coverage on the test set.

#### Gene-level correlation

We aggregated exon coverage to a gene expression value for each track for each gene in the test set. Then for each track, we computed the correlation between predicted and observed gene expression on the test set genes.

#### Bin-level specificity

We constructed observed and predicted track-by-bin coverage matrices on the test set. For each of the two matrices, we performed quantile normalization to account for sequencing depth across tracks and computed the log fold change difference of each track versus the average across all tracks at the bin level. For each track, we computed the correlation between the predicted and observed log fold change as the bin-level specificity score.

#### Gene-level specificity

We aggregated exon coverage to a gene expression value for each track for each gene in the test set. For each of the two matrices, we performed quantile normalization to account for sequencing depth across tracks and computed the log fold change difference of each track versus the average expression profile across all tracks. For each track, we computed the correlation between the predicted and observed log fold change as the gene-level specificity score.

### 5.6 Attribution methods

#### Gene Saliency Score

We applied the Borzoi ensemble model across 4 folds to compute the saliency of aggregated RNA-seq coverage across exon bins with respect to a 524 kb input sequence for any gene of interest, which we refer to as the **gene saliency score**. Since the Borzoi model has greater gradients closer to the gene’s promoter and exons, we sought to enhance signal detection in more distal regions. To achieve this, we calculated a **normalized gene saliency score** by first applying local smoothing to the saliency score using a 1D Gaussian filter (sigma = 5), followed by z-score normalization based on the mean and standard deviation within a 10 kb window.

#### Differential Saliency Score (DSS)

To compare treatment and control RNA-seq tracks, we used the Borzoi ensemble model to first compute the gene saliency score for a gene of interest. We then normalized the two saliency vectors by matching the 20% quantile of the absolute saliency, and took the difference between the two vectors (treatment minus control). This gives us a differential saliency score vector, which quantifies contribution to differential coverage over the gene in treatment versus control.

#### Normalized Differential Saliency Score (NDSS)

To improve the signal in more distal regions, we performed local smoothing on the differential saliency score with gaussian filter1d(sigma=5) and then performed z-score normalization using the mean and standard deviation within a 10 kb window.

#### Motif Instance Score

We used an existing database of all the motifs in the JASPAR2022

CORE vertebrates redundant pfms mapped to the hg38 genome from https://github.com/wassermanlab/ JASPAR-UCSC-tracks. For every motif instance with p<1e-4 within the 524k context window of our gene of interest, we computed the mean DSS, mean NDSS, and PWM-gradient correlation. Motif instances in non-TSS exonic regions were excluded because Borzoi’s high attribution in these areas reflects coding gene modifications rather than transcriptional regulatory function. Motif instances with abs(NDSS)>1.96 and PWM-gradient correlation significantly greater than permutation (*p_adj_ <* 0.01) were considered significant motif instances that contribute to differential regulation of the gene. The motif instance score was computed by taking the average of NDSS in the motif window.

### 5.7 Hayflick benchmark

#### 5.7.1 WI-38 cCRE classification

We downloaded a comprehensive human cCRE atlas compiled by ENCODE from https://screen.wenglab.org, which contains a total of 1,063,878 cCREs [22]. To classify peaks, we defined ENCODE cCREs overlapping with the WI-38 ATAC atlas, generated by Chan et al. [5], as positives (i.e., peaks accessible in WI-38). Conversely, ENCODE cCREs that do not overlap with the WI-38 ATAC atlas were classified as negatives (i.e., peaks not accessible in WI-38).

For each gene of interest, we computed normalized gene saliency scores based on SEN TP1 RNA-seq coverage with respect to the input DNA sequence. We prioritized ENCODE cCREs within a 524 kb window around the gene center by taking the maximum absolute value of the normalized gene saliency scores within each cCRE. Using this approach, we evaluated the classification performance distinguishing WI-38-accessible peaks from inaccessible ones. We performed this classification on both all ENCODE cCREs and enhancer cCREs only (excluding cCREs located less than 1kb from TSS).

#### 5.7.2 Differential TF activity analysis

##### ATAC-based enrichment

As ground truth, we used the open chromatin profiles assayed at TP7 and TP1 in combination with the RNA-seq data to identify TFs responsible for differential gene regulation in replicative senescence (SEN TP7 vs TP1). (1) We performed FIMO motif search on the WI-38 ATAC-seq atlas for all motifs in the JASPAR2022 CORE vertebrates redundant pfms collection to derive a binary peak-by-motif matrix. (2) Peaks with significantly increased accessibility (*p_adj_ <* 0.05 and logFC>0) located near upregulated genes (*p_adj_ <* 0.01, logFC>1, peak-gene distance<50kb) were considered up-regulating DARs. (3) Peaks with significantly decreased accessibility (*p_adj_ <* 0.05 and logFC<0) located near down-regulated genes (*p_adj_ <* 0.01, logFC<-1, peak-gene distance<50kb) are considered down-regulating DARs. (4) For each TF, we performed hypergeometric tests to assess motif enrichment in up- and down-regulating DARs, reporting the odds ratio for up-regulating DARs and the negative odds ratio for down-regulating DARs. Finally, for each TF, we reported the odds ratio with the largest absolute value, indicating the stronger direction of regulation.

##### RNA-based promoter enrichment

Without ATAC-seq information, we performed a similar motif enrichment analysis focusing on promoter regions only. (1) We performed FIMO motif search on the DEG promoters (2kb window from TSS) for all motifs in the JASPAR2022 database to derive a binary promoter-by-motif matrix. (2) For each TF, we performed hypergeometric tests to assess motif enrichment in promoters of up- and down-regulating genes, reporting the odds ratio for up-regulating promoters and the negative odds ratio for down-regulating promoters. Finally, for each TF, we reported the odds ratio with the largest absolute value, indicating the stronger direction of regulation.

##### RNA-based Borzoi prediction

Alternatively, without information on ATAC, we used the Borzoi model to identify key TFs that drive differential gene regulation in the TP7 vs. TP1 RNA-seq comparison. For each TF motif in the JASPAR database, we first computeed NDSS scores (TP7 vs. TP1) and identified significant motif instances around the top 5000 differentially expressed genes as described above. We then summarized TF contribution to up-regulated genes by taking the mean NDSS scores across all significant motif instances near these genes and computed a contribution to down-regulated DEGs in the same way. We determined the final direction of the TF motif activity by comparing the absolute value of the two scores.

### 5.8 TFKD benchmark

For each TFKD experiment, we evaluated the efficacy of CRISPRi/RNAi knockdown by the logFC of targeted TF mRNA level. Experiments with targeted TF logFC<-2, -2≤ logFC<-1, -1≤logFC<0, and 0≤logFC were categorized into strong, medium, weak and no effect groups.

There are 28 TFKD experiments in the strong/medium groups where the TF’s motif also exists in the JASPAR2022 CORE vertebrates redundant pfms database. For each of these experiments and corresponding control RNA-seq, we identified a set of down-regulated genes (down-DEGs) using the gene TPM quantification. We selected up to 1000 down-DEGs with TPM>1 sorted by logFC. Then, we evaluated whether the Borzoi model could recapitulate the reduced activity of the targeted TF in the down-DEGs and compared against a simple TSS motif enrichment approach.

#### 5.8.1 TSS motif enrichment

As a baseline, we implemented the SEA tool in the MEME Suite for TSS motif enrichment analysis [2]. For each TF knockdown RNA-seq experiment, we used 2kb sequence centered around the TSS of the down-DEGs as the positive set and 2kb sequence centered around the TSS of 1000 random genes (excluding up-DEGs and down-DEGs) as the negative set. We ran SEA with JASPAR2022 CORE vertebrates redundant pfms motif database and arguments ’–no-pgc –pvalue –thres 1’ to report statistics for all motifs.

For each of the 28 TFKD experiments, we ranked all TFs by negative log10 p-value and asked whether the targeted TF was ranked among the top 10, 20, 50 and 100.

#### 5.8.2 Borzoi TF activity analysis

For each perturbation experiment and the corresponding control, we first computed the differential saliency score (DSS) and normalized differential saliency score (NDSS) vectors for each down-regulated DEG. We then computed motif instance scores and identified significant motif instances contributing to the down-regulated DEGs using the method described above.

For a given TFKD experiment, to identify top TFs regulating the down-DEGs, we only considered significant motif instances with negative DSS scores, as these instances contribute to the down-regulation of genes. For each TF, we summed up DSS motif instance scores in promoters (2kb from TSS), enhancers (distal), or combined, to study whether their contribution is proximal or distal.

For each of the 28 TFKD experiments, we ranked all TFs by Borzoi TF activity score (promoter, enhancer, or combined) and asked whether the targeted TF was ranked among the top 10, 20, 50 and 100.

#### 5.8.3 ENCODE TF ChIP-seq

For selected TFs, we also downloaded the signal p-value BigWig track and IDR-thresholded narrowPeak files from ENCODE K562 ChIP-seq datasets for evaluation (ENCSR740NPG for BACH1, ENCSR028UIU for ATF3 and ENCSR000DJX for JUND).

### 5.9 PBMC benchmark

#### 5.9.1 Differential TF activity analysis

##### Borzoi derived TF activity scores

To infer TF activity in each cell type using Borzoi with only the scRNA-seq modality, we followed the same approach described in previous sections. Specifically, we first computed NDSS scores for motif instances located near the top 500 genes that are differentially up-regulated in the target cell type versus all others, across all TFs in the JASPAR database. TF activity was then summarized by averaging NDSS scores across significant motif instances near these up-regulated genes, followed by z-score normalization.

##### scBasset derived TF activity scores

We trained an scBasset model on the scATAC modality of 10x ARC PBMC. To infer TF activity in each cell type using scBasset, we scored all JASPAR motifs at single-cell resolution using the motif insertion approach as described in the original publication [34]. For each target cell type, we then performed a two-sample t-test comparing motif scores between cells of interest and all other cells, using the resulting test statistic as a measure of differential TF activity.

#### 5.9.2 Onek1k eQTL effect size prediction

We downloaded summary statistics and fine-mapping results from the Onek1k dataset [32]. For each cell type of interest, we selected all fine-mapped variants with a posterior inclusion probability (PIP)>0.9 and scored them using the corresponding pseudobulk track from the transferred Borzoi model. We then computed the Pearson correlation between the measured and Borzoi-predicted effect sizes.

## 6 Data availability

We used only public datasets in this study. Pre-trained full Borzoi model is obtained from Linder at al. [19]. The GTEx dataset contains 89 unstranded RNA-seq coverage tracks from GTEx whole-tissue samples [7], uniformly processed by the recount3 project and later subsampled by Linder et al. [19, 31]. The Hayflick dataset is obtained from Chan et al. [5]. ENCODE TFKD dataset is downloaded from ENCODE portal (https://www.encodeproject.org/), with accession number available in Supplemetary Table S7. Variant-FlowFISH dataset is obtained from Martyn & Montgomery et al. [21]. The 10x PBMC RNA+ATAC multiome dataset was downloaded from https://support.10xgenomics.com/single-cell-multiome-atac-gex/datasets/1.0.0/pbmc_granulocyte_sorted_10k.

## 7 Code availability

Code for efficient Borzoi transfer is integrated into the Baskerville codebase: https://github.com/calico/baskerville. An example tutorial for Borzoi transfer is available at https://github.com/calico/baskerville/blob/main/docs/transfer/transfer.md.

## 8 Acknowledgements

This work was funded by Calico Life Sciences. The funder had no role in study design, data collection, or analysis. Publication of the manuscript was approved after an internal scientific review process. We thank Ian Holmes, Divyanshi Srivastava, Fanny Huang, Anya Korsakova, Benjamin Auerbach, David Wang, Luong Ruiz, Jun Xu, Michael Closser, Wenjun Kong, and Alexis Leigh Krup for helpful discussions and valuable feedback.

## 10 Supplementary Figures

**Supplementary Figure S1:**
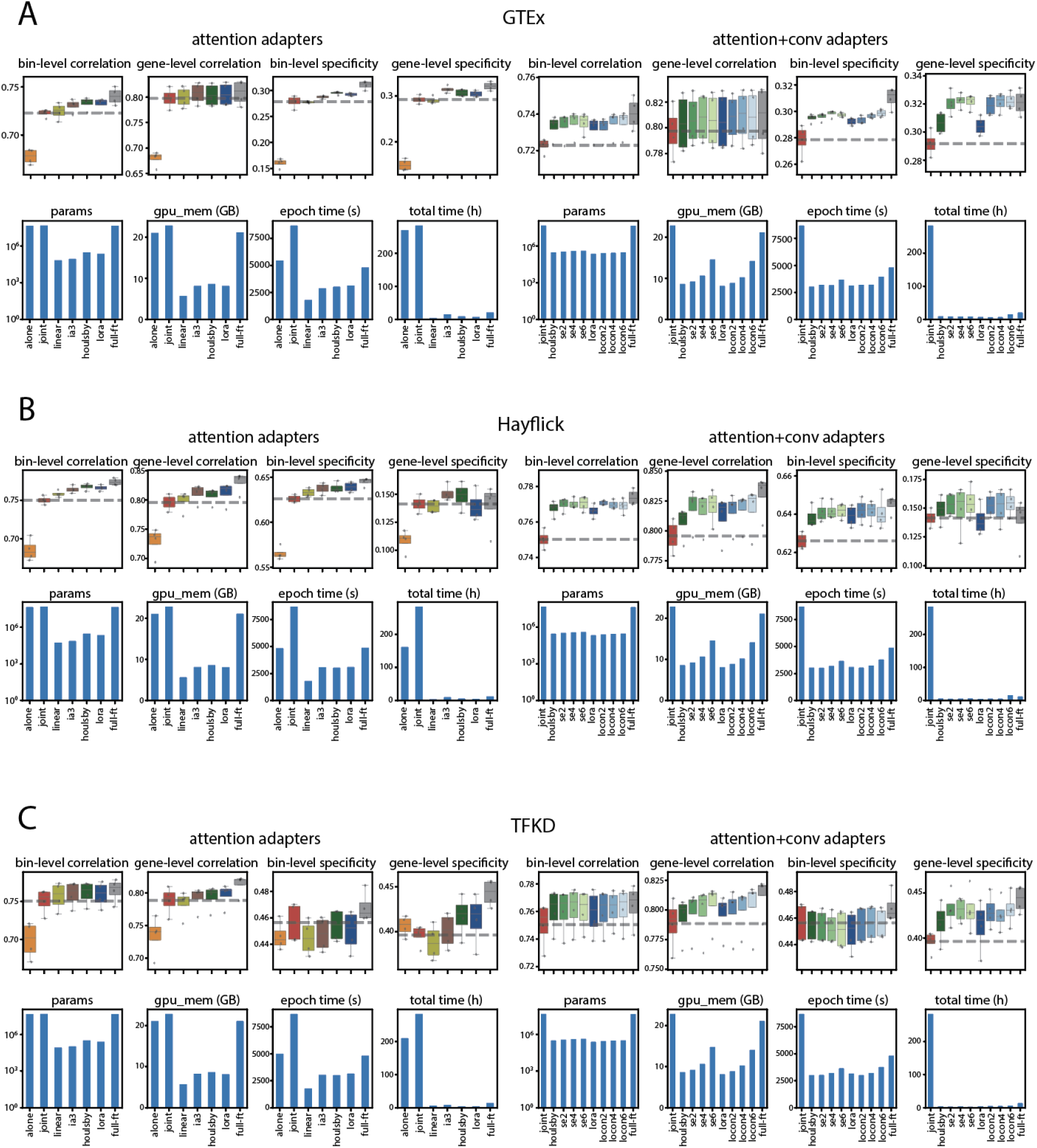
**(A)** Transfer performance of the Borzoi-lite-no-GTEx model using IA3, Houlsby, and LoRA adapters compared to baseline methods on GTEx test sequences (left). Benchmark of adapter insertion strategies—Locon and Squeeze-Excitation—applied to 2, 4, or 6 convolutional layers on top of LoRA or Houlsby (right). Baseline methods include training from scratch on new data only (alone), joint training with pre-training data (joint), linear probing (linear), and full fine-tuning (full-ft). Details on evaluation metrics are provided in the Methods section. Transfer benchmarks for the Borzoi-lite model on the Hayflick and TFKD datasets are shown in **(B)** and **(C)**, respectively.

**Supplementary Figure S2:**
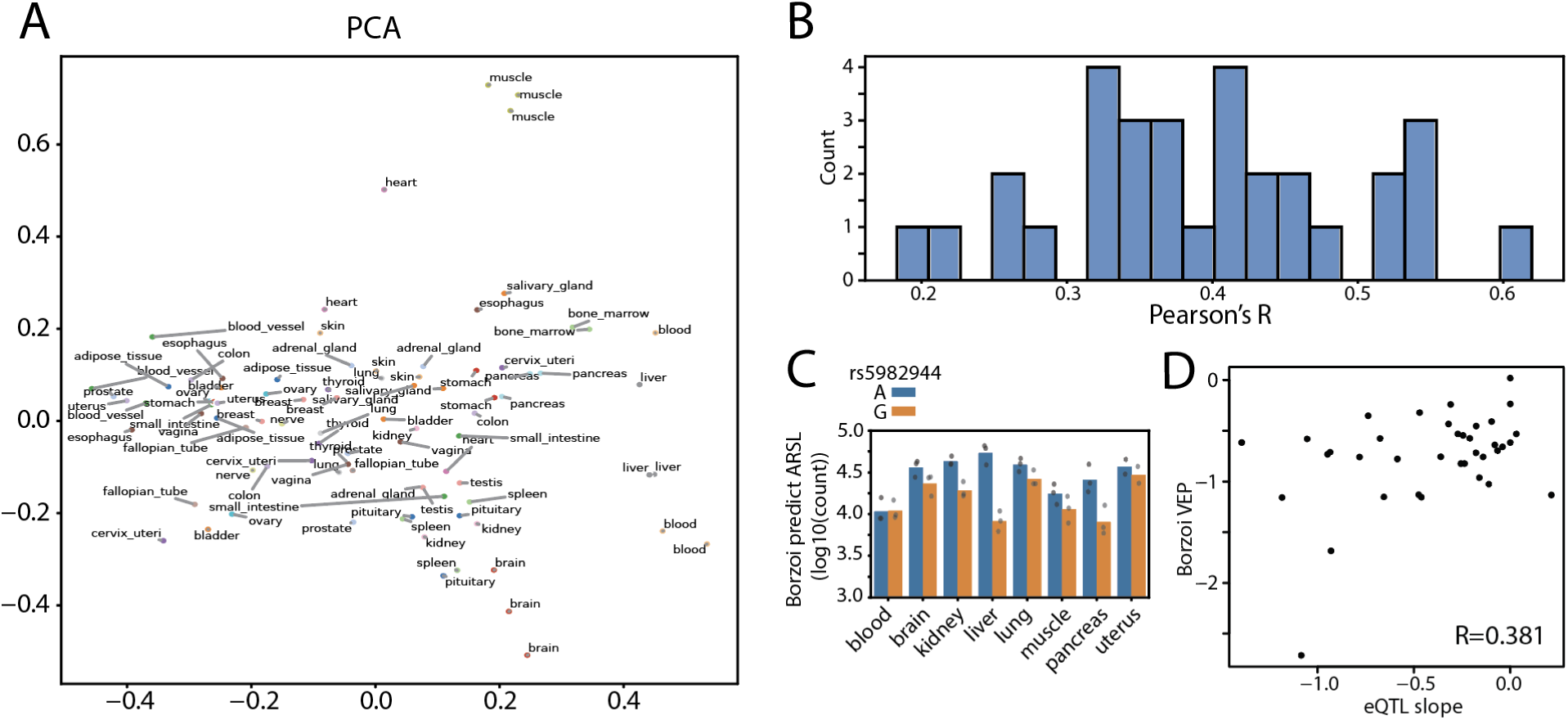
**(A)** PCA embeddings of the learned task representations from the final model layer parameters for all of 89 GTEx tracks representing 31 tissue types. **(B)** Histogram of Pearson correlation coefficients between observed and predicted gene expression logFC on test set, comparing each tissue of interest to all other tissues. **(C)** Borzoi-predicted expression of *ARSL* for the A and G alleles at rs5982944. Points represent prediction for individual GTEx track. **(D)** Borzoi prediction versus measure eQTL effect size fo rs5982944 across all tissue.

**Supplementary Figure S3:**
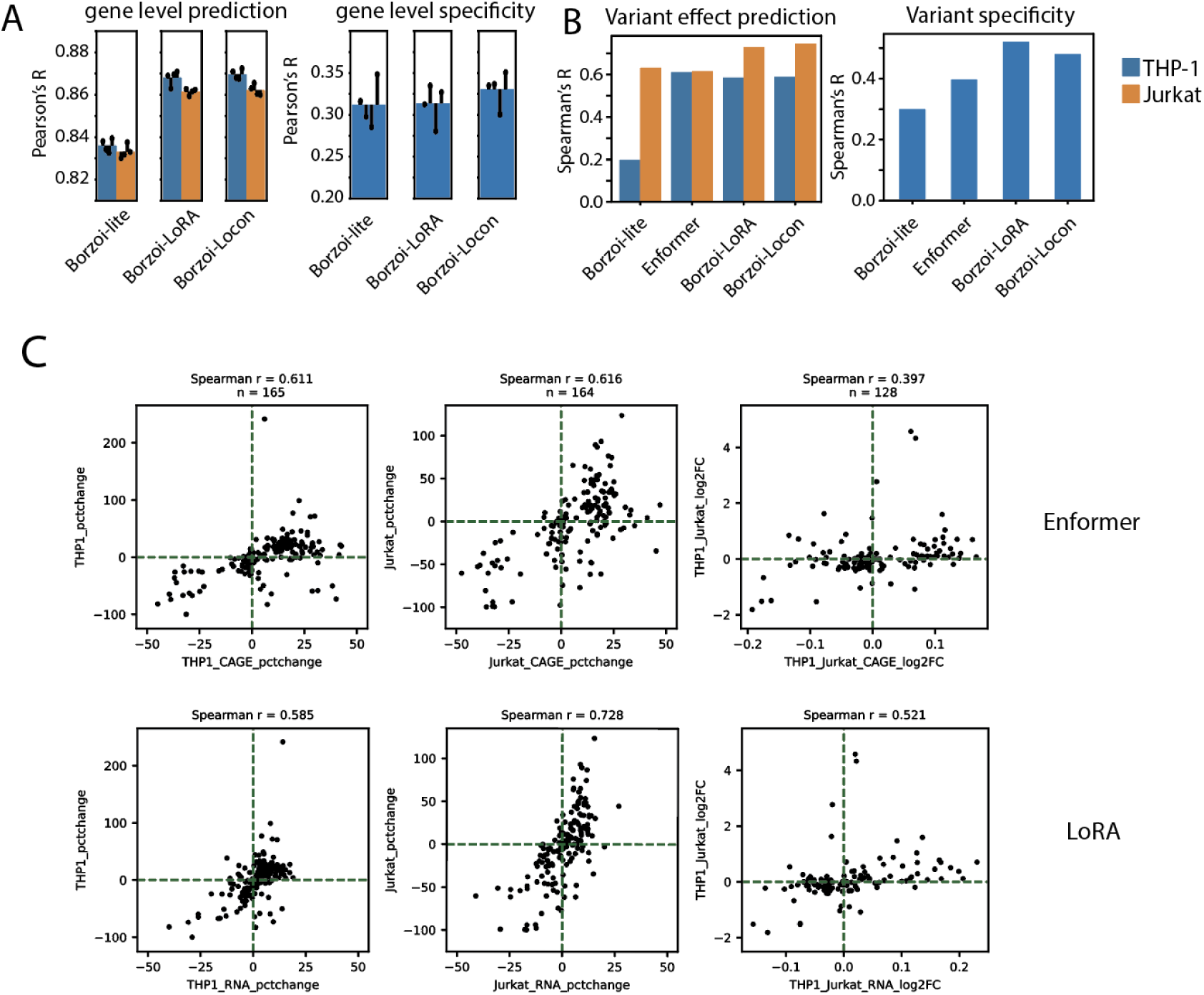
**(A)** Comparison of Borzoi model transferred to variant Flowfish dataset using LoRA, Locon4 or training from scratch a Borzoi-lite model in gene-level correlation (left), and gene-level specificity (right) on test genes. **(B)** Performance comparison of variant effect prediction for THP-1 and Jurkat, quantified by Spearman’s correlation between predicted and measured changes in PPIF expression (left), and cell-type specificity, quantified by Spearman’s correlation between predicted and measured logFC differences between THP-1 and Jurkat (right). Comparisons include Borzoi-lite models trained from scratch, Enformer CAGE tracks of matched cell types, Borzoi with LoRA transfer, and Borzoi with Locon4 transfer. **(C)** Comparison of measured versus Enformer(top)/LoRA(bottom) predicted variant effects on PPIF expression for variants in THP-1 (left, quantified by percentage change in PPIF expression), Jurkat (middle, quantified by percentage change in PPIF expression), and specificity (right, quantified by logFC difference between THP-1 and Jurkat). Spearman’s correlation between predictions and measurements is shown in the top left of each panel.

**Supplementary Figure S4:**
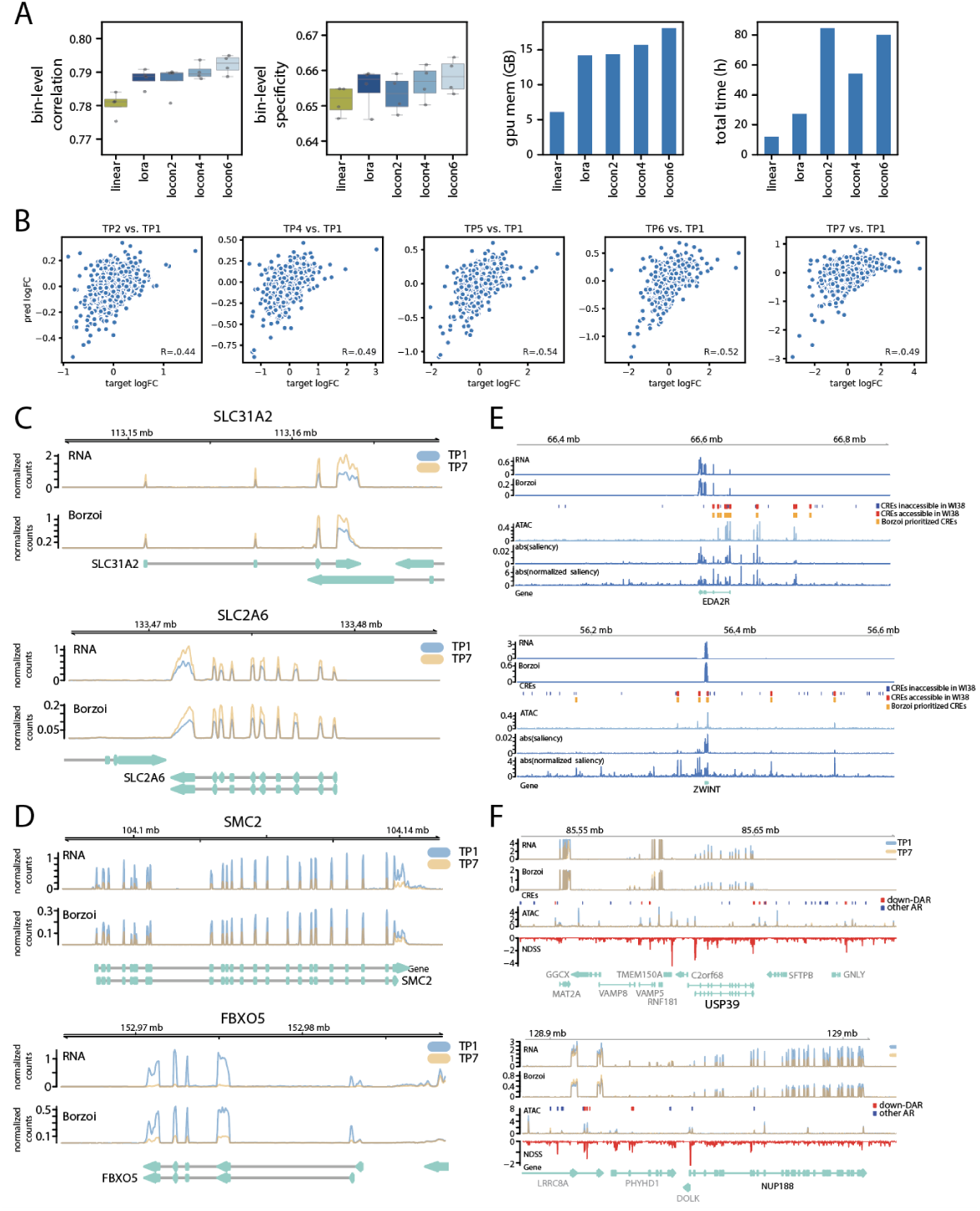
**(A)** Full Borzoi transfer performance on Hayflick dataset measured by bin-level correlation and specificity on test sequences. We compared between LoRA, Locon with inserted to 2, 4 and 6 convolutional layers, and linear probing. We also compare the maximum GPU memory usage and total run time. **(B)** Correlation between Borzoi prediction and observed held-out gene expression logFC during senescence (from left to right: TP2, 4, 5, 6, 7 vs. TP1). TP3 is excluded due to low data quality [5]. **(C-D)** Pedicted and observed tracks around senescence-induced genes *SLC31A2* and *SLC2A6* (C), and senescence-repressed genes *SMC2* and *FBXO5* in the test set. RNA: observed RNA-seq for TP1 (control) and TP7 (senescent) samples. Borzoi: Borzoi predicted RNA-seq for TP1 and TP7. **(E)** Two examples of normalized gradient around test genes EDA2R and ZWINT overlap with accessible chromatin. RNA: observed RNA coverage at TP1. Borzoi: predicted RNA coverage at TP1. CREs: All ENCODE CREs in 500kb window are labeled in blue, those overlap WI38 ATAC peaks are labeled in red. Those prioritized by Borzoi saliency are labled in orange. ATAC: observed ATAC track at TP1. abs(saliency): absolute value of saliency of aggregated RNA-seq coverage across exon bins with respect to a 524 kb input sequence for gene of interest. abs(normalized saliency): absolute value of the locally smoothed saliency (Methods). **(F)** Pedicted and observed tracks around senescence-repressed genes USP39 and NUP188. RNA: observed RNA-seq for TP1 (control) and TP7 (senescent) samples. Borzoi: Borzoi predicted RNA-seq for TP1 and TP7. ATAC: observed ATAC-seq for TP1 and TP7, with differentially accessible regions (DAR) labeled in red, and other accessible regions (AR) in green. NDSS: Normalized Differential Saliency Scores (NDSS) comparing the saliency between TP1 and TP7 (Methods).

**Supplementary Figure S5:**
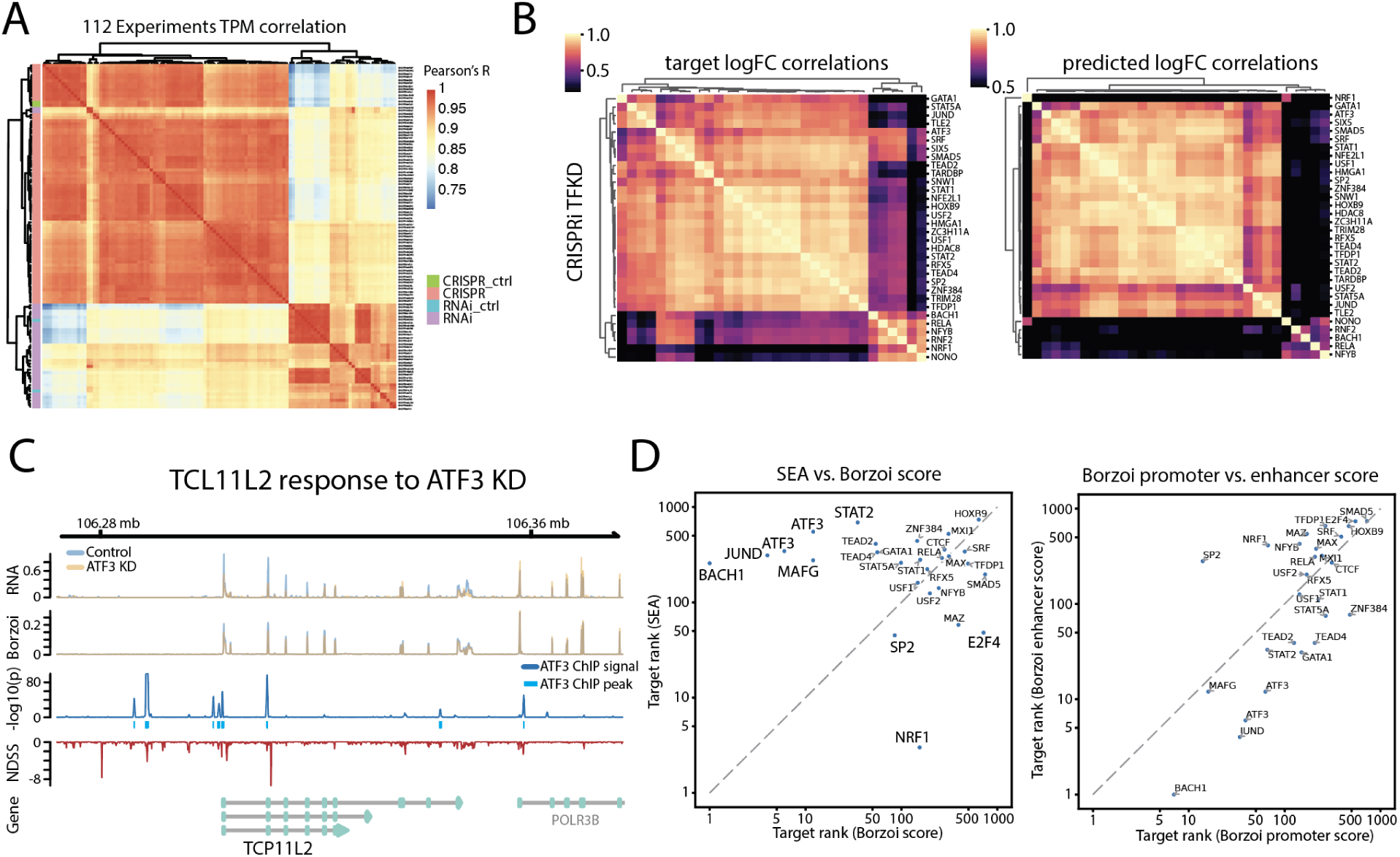
**(A)** Gene expression correlation across all 112 experiments reveals clear batch effect between CRISPR-based and RNAi based experiments. **(B)** Correlation heatmaps of observed (left) and predicted (right) expression change (quantile normalized, perturbed versus control logFC) on test genes across CRISPR experiments. **(C)** Predicted and observed tracks around a repressed gene *TCP11L2* after ATF3 CRISPRi knockdown. RNA: observed average RNA-seq tracks for control and ATF3 KD samples. Borzoi: Borzoi predicted RNA-seq tracks for control and ATF3 KD samples. ChIP: ATF3 ChIP-seq signal p-value track in K562. IDR thresholded peaks are shown in light blue. NDSS: Normalized Differential Saliency Scores (NDSS) comparing the saliency of *TCP11L2* between ATF3 KD and control sample (Methods). Gene: Gene annotation track. **(D)** Left, for 28 knockdown experiments where the target TF has an existing motif in the JASPAR2022 database, we reported the ranks of the perturbed TFs in each experiment using Borzoi TF score, or SEA based on TSS enrichment. Right, we reported the ranks of the perturbed TFs in each experiment using Borzoi TF scores from promoter only, or enhancer only.

**Supplementary Figure S6:**
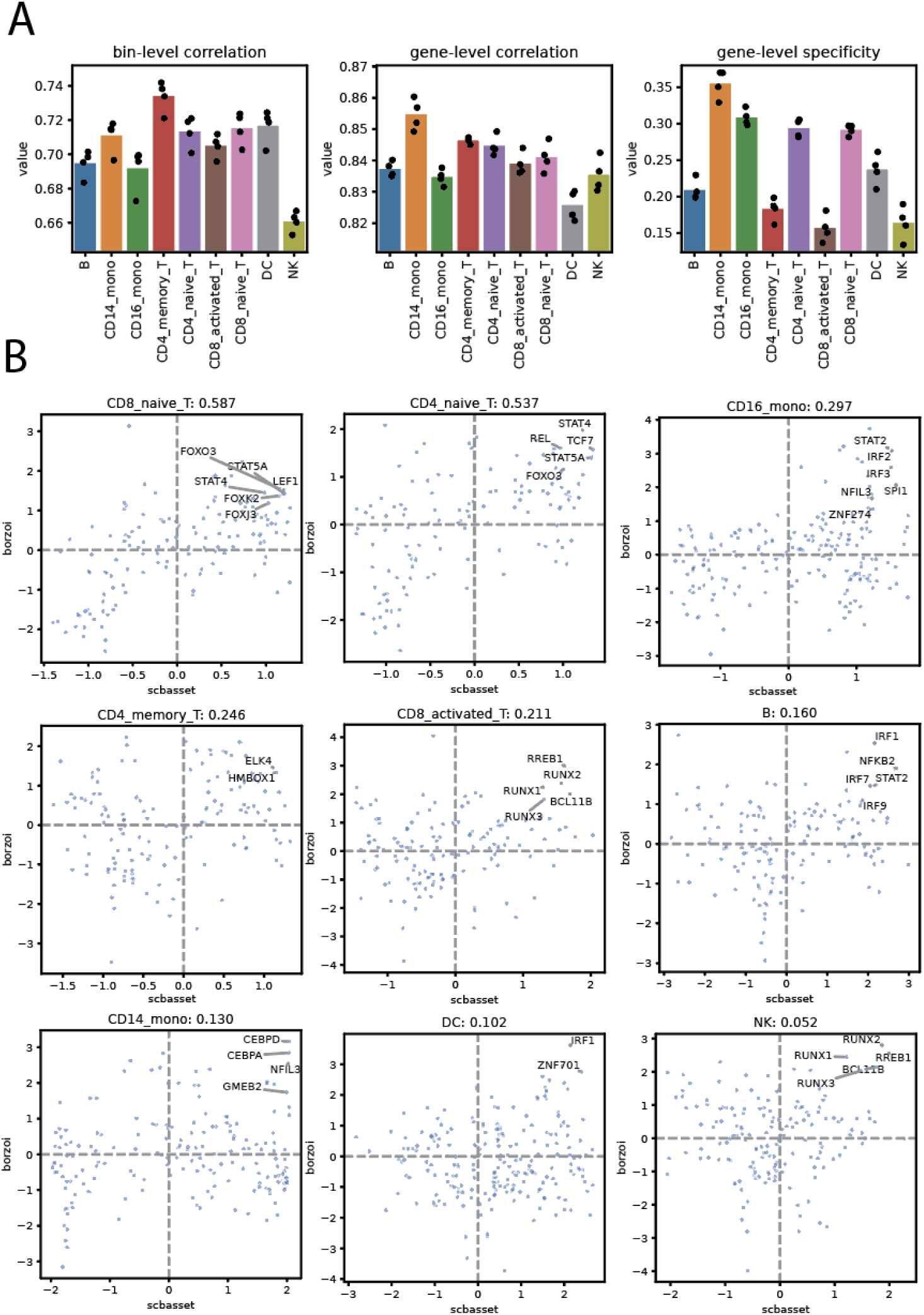
**(A)** Test set performance of Locon4 Borzoi transfer model on PBMC dataset. Details of metrics are in the Methods section. **(B)** Comparison between TF activity inferred from scBasset using scATAC modality (x-axis) versus activity inferred from Borzoi using scRNA modality (y-axis) for all cell types.

**Supplementary Figure S7:**
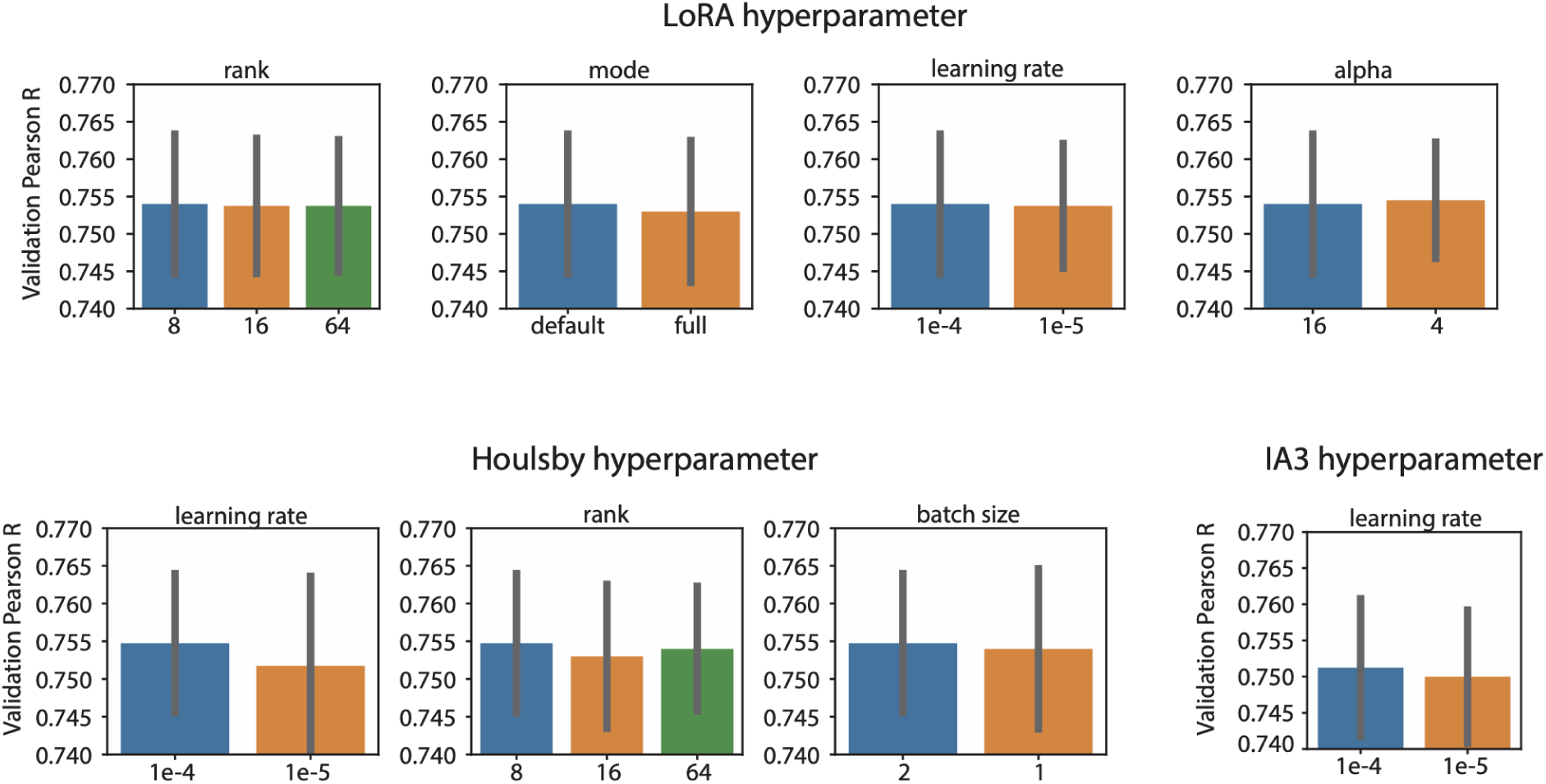
**(A)** Test set performance of Locon4 Borzoi transfer model on PBMC dataset. Details of metrics are in the Methods section. **(B)** Comparison between TF activity inferred from scBasset using scATAC modality (x-axis) versus activity inferred from Borzoi using scRNA modality (y-axis) for all cell types.

